# Lipotype acquisition during neural development *in vivo* is not recapitulated in stem cell-derived neurons

**DOI:** 10.1101/2022.08.31.505694

**Authors:** Anusha B. Gopalan, Lisa van Uden, Richard R. Sprenger, Nadine Fernandez-Novel Marx, Helle Bogetofte, Pierre Neveu, Morten Meyer, Kyung-Min Noh, Alba Diz-Muñoz, Christer S. Ejsing

## Abstract

During development, different tissues acquire distinct lipotypes that are coupled to tissue function and homeostasis. In the brain, where complex membrane trafficking systems are required for neural function, specific glycerophospholipids, sphingolipids, and cholesterol are highly abundant, and defective lipid metabolism is associated with abnormal neural development and neurodegenerative disease. Notably, the production of tissue-specific lipotypes requires appropriate programming of the underlying lipid metabolic machinery, but when and how this occurs is unclear. To address this, we used high-resolution mass spectrometry-based (MS^ALL^) lipidomics to perform a quantitative and comprehensive analysis of mouse brain development covering early embryonic and postnatal stages. We discovered a distinct bifurcation in the establishment of the neural lipotype, whereby the canonical brain lipid biomarkers 22:6-glycerophospholipids and 18:0-sphingolipids begin to be produced *in utero*, whereas cholesterol attains its characteristic high levels after birth. In contrast, when profiling rodent and human stem cell-derived neurons, we observed that these do not acquire a brain lipotype *per se*. However, upon probing the lipid metabolic wiring by supplementing brain lipid precursors, we found that the stem cell-derived neurons were partially able to establish a brain-like lipotype, demonstrating that the cells are partially metabolically committed. Altogether, our report provides an extensive lipidomic resource for brain development and highlights a potential challenge in using stem cell-derived neurons for mechanistic studies of lipid biochemistry, membrane biology and biophysics that can be mitigated by further optimizing *in vitro* differentiation protocols.

**Significance Statement:** We report an extensive time-resolved resource of lipid molecule abundances across mouse brain development, starting as early as 10 days post-fertilization. The resource reveals a bifurcation in the establishment of the neural lipotype where the canonical 22:6-glycerophospholipid and 18:0-sphingolipid biomarkers are attained *in utero*, whereas cholesterol is attained after birth. Furthermore, we uncover that the neural lipotype is not established in rodent and human stem cell-derived neurons *in vitro*.

## Introduction

Lipids are a diverse category of biomolecules that constitute and functionalize membranes, serve as energy reservoirs, and function as signaling molecules. While different tissues and cell types produce and maintain distinct characteristic lipid compositions, termed lipotypes (Hicks et al., 2006; Harayama et al., 2014; Capolupo et al., 2022), how they are acquired during the course of development is largely unknown. Furthermore, the functional implications of maintaining a distinct lipotype are not understood, although the existence of tissue-specific lipid homeostasis across organisms points to conserved mechanisms governing critical functions (Yamashita et al., 2014; Bozek et al., 2015).

Lipotype acquisition and homeostasis are of particular interest in neuroscience due to the enrichment of distinct lipid molecules in the brain, which is functionally coupled to the sophisticated membrane trafficking system of neural cell types (Davletov & Montecucco, 2010; Puchkov & Haucke, 2013; Lauwers et al., 2016; Ingólfsson et al., 2017). Studies have previously reported severe impairments of neuronal development in mice lacking the lipid binding protein SCAP (Verheijen et al., 2009) as well as the lipid metabolic enzymes CerS2 (Imgrund et al., 2009) and DAGL (Gao et al., 2010), which are part of the cholesterol biosynthesis, ceramide (Cer) biosynthesis and diacyl glycerol breakdown pathways, respectively. Defects in sphingolipid metabolism have also been associated with neural tube defects in embryos (Stevens & Tang, 1997; Missmer et al., 2006), the degeneration of motor neurons (Bejaoui et al., 2001; Dawkins et al., 2001) and myelination defects (Fewou et al., 2005). Moreover, the loss of ether-linked phosphatidylethanolamines (PE O-, i.e., plasmalogens) and sulfatides have been linked to age-related neurodegeneration in diseases such as Alzheimer’s (Han et al., 2001, 2002; Tu et al., 2018). Biochemical studies have also shed light on the various roles of sphingomyelin (SM) and the ganglioside GM1, and potential lipid signaling molecules such as retinoids, terpenoids, steroids, and eicosanoids, in triggering differentiation programs in neuronal stem cells (Bieberich, 2013). Furthermore, studies performed *in vitro* (Cao et al., 2009; Pinot et al., 2014) as well as *in vivo* (Janssen et al., 2015) have demonstrated that docosahexaenoic acid (DHA, 22:6), found in membrane glycerophospholipids, is important for neurogenesis and the formation of synapses (Salem et al., 2001; Innis, 2007). Thus, lipids have been identified as key players across neurological development, physiology and disease.

Decades of research on brain lipids of adult animals has established a characteristic high level of 22:6-containing glycerophospholipids, 18:0-containing sphingolipids and cholesterol as canonical brain lipid biomarkers (O’Brien et al., 1964; O’Brien & Sampson, 1965; Fitzner et al., 2020). Since these lipids are membrane constituents, it is possible that their levels are tightly coupled to membrane trafficking events and activities of cell surface receptors. From a medical perspective, studying their role can further our understanding of the developmental deficits and diseases related to lipid metabolism in neurons, and also inform procedures in regenerative medicine involving the differentiation of stem cells.

Recent improvements in lipid mass spectrometry now allows in-depth quantitative lipidomics of lipid extracts, from minute amounts of tissues and cells, to obtain information on the molecular identity of thousands of lipid molecules, down to lipid class and individual fatty acyl chains (Almeida et al., 2014). Recent work has built on characterizing the lipidomes of adult mouse brain tissue as well as its constituent cell types (Fitzner et al., 2020). The canonical brain lipid biomarkers discussed here are found to be enriched across distinct neural cell types, demonstrating a tissue-specific, rather than a cell-type-specific lipotype in the brain. This is further supported by the overall lipidome similarity of subcellular postsynaptic densities isolated from rat neurons to brain tissue (Tulodziecka et al., 2016).

To shed light on how the brain lipotype is acquired during development, we performed a lipidomic analysis of brain tissue over the course of early brain development in mice, starting from the embryonic stage of E10.5 where the brain region first becomes accessible for dissection, up to the postnatal stage P21. We find that high levels of the canonical brain lipid biomarkers 22:6-glycerophospholipids and 18:0-sphingolipids begin to be established already at the embryonic stage *in utero*, coinciding with extensive neurogenesis (E10.5-E15.5) (Caviness, 1982; Finlay & Darlington, 1995), while the increase in cholesterol occurs postnatally. To investigate the interplay between the establishment of lipid metabolic programs and the acquisition of neuronal fate, we turned to stem cell-derived neurons. Unlike neural cells *in vivo*, we found that neither mouse embryonic stem cells (mESCs) nor human induced pluripotent stem cells (hiPSCs) undergo lipidomic remodeling during neuronal differentiation. To mitigate this deficiency, we supplemented the culture medium with lipid precursors required to synthetize key brain lipid biomarkers during differentiation. This supplementation partially established the levels of the canonical brain lipid markers, demonstrating that the lipid metabolic machinery to synthetize 22:6-glycerophospholipids is set up in stem cell-derived neurons, while that responsible for synthesis of 18:0-sphingolipids is not. Taken together, our findings show that early mouse brain development coincides with extensive lipidome remodeling, starting already at the embryonic stage and that *in vitro* stem cells are only partially metabolically committed.

## Results

### Lipotype acquisition during early mouse brain development

To identify lipid hallmarks associated with neural development, we compiled a comprehensive lipidomic resource of mouse brain development. To this end, we micro-dissected the whole mouse brain region at the developmental stage E10.5, as well as excised cerebral hemispheres from E15.5, P2 and P21 mice (**Fig. 1A**). These biopsies were analyzed by high-resolution MS^ALL^ lipidomics (Almeida et al., 2014; Sprenger et al., 2021). Overall, this analysis identified and quantified 1488 lipid molecules encompassing 26 lipid classes (**Table S1**).

**Figure 1.**
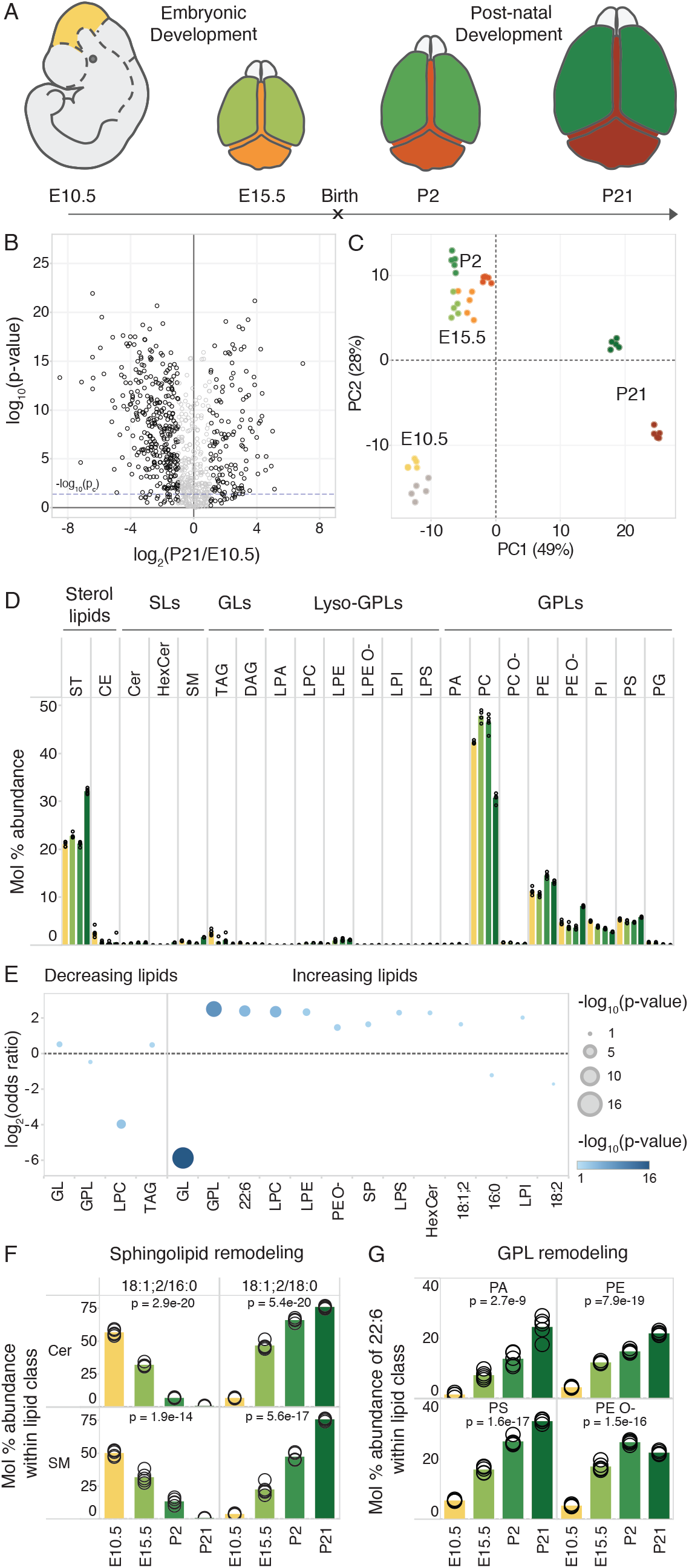
Lipotype acquisition during brain development. **A)** Mouse brain tissue was collected at the indicated time points and analyzed by in-depth MS^ALL^ lipidomics. The colors denote distinct brain regions that were sampled at each developmental stage. **B)** Volcano plot of molecular lipid species. The fold-change of each lipid within its lipid class is calculated between the E10.5 whole brain and P21 cerebral hemisphere. **C)** Principal component analysis of lipid abundances across samples, colored as per the schematic in A. **D)** Profile (mol%) of different lipid classes across developmental time. Bars representing the mean value for each sample group are colored in accordance with panel A. **E)** Lipid feature ENrichment Analysis (LENA) of features among decreasing and increasing lipids, where -log_2_ (odds ratio) is plotted in the y-axis and the color and size correspond to the -log_10_(p-value). **F)** Profile of the most abundant Cer and SM species across samples, containing an 18:1;2 sphingoid chain and either a 16:0 or 18:0 acyl chain. **G)** Profile of 22:6 (DHA) among fatty acyls constituting phosphatidic acid (PA), PE, PE O- and PS species. Data was collected on n = 5 replicates. p-values were obtained using a one-way ANOVA test and changes were considered significant if p ≤ p_c_ = 0.04, as determined by the Benjamini-Hochberg procedure.

Statistical analysis revealed that 451 distinct lipid molecules in the cerebral hemispheres (30% of detected lipids) were significantly changed in abundance across the developmental timeline with a two-fold or greater difference between E10.5 and P21 (ANOVA followed by the Benjamini-Hochberg procedure, p ≤ p_c_ = 0.04) (**Fig. 1B**), of which 133 (9%) and 318 (21%) lipids increased and decreased, respectively. Principal component analysis showed a clustering of samples according to time-point and tissue, with only a weak distinction between the E10.5 brain tissue and the rest of the embryo (**Fig. 1C, S2, S3**). Similarly, the distinction between cerebral hemispheres and the rest of the brain tissues was not pronounced at E15.5 but clearly increased over the course of development as each brain region acquired a more specialized lipotype.

Assessing the bulk abundance of lipid classes in brain tissue across development, we observed a 1.5-fold increase in cholesterol and a 2-fold increase in PE O-as well as a corresponding decrease in phosphatidylcholines (PC) in the postnatal phase between P2 and P21. Low levels of cholesterol esters (CE) and triacylglycerol (TAG) present at E10.5 were further diminished thereafter. The total level of phosphatidylinositols (PI), precursors of signaling phosphoinositides, gradually decreased in abundance throughout development (**Fig. 1D**).

To systematically examine whether the lipids increasing in abundance have common molecular traits, we carried out a Lipid feature ENrichment Analysis (LENA) (Sprenger et al., 2021), akin to gene ontology analysis for gene transcripts and proteins. This analysis demonstrated that membrane glycerophospholipids (GPL, p = 8·10^−10^) and sphingolipids (SP, p = 7·10^−3^) are enriched among the increasing lipid molecules whereas storage glycerolipids (GL, p = 1·10^−16^) are depleted from this pool. Moreover, we found that the lysolipids LPC, LPE, LPS and LPI are enriched in the pool of increasing lipids, in addition to hexosylceramides (HexCer) and PE O-lipids. The polyunsaturated structural attribute 22:6 (p = 6·10^−6^), corresponding to DHA, is significantly enriched among the increasing lipids, as are sphingolipids with a sphingosine 18:1;2 chain (p = 2·10^−2^) (**Fig. 1E**). Inspecting the molecular timelines of the canonical brain lipid biomarkers showed that the sphingolipids Cer 18:1;2/18:0 and SM 18:1;2/18:0 as well as 22:6-containing phosphatidylserine (PS), PE and PE O-species already reach their expected high levels within the early developmental period studied here (**Fig. 1F and 1G**).

For all time-points, we also subjected the remainder of the brain (after removal of the cerebral hemispheres) to lipidomic analysis and found similar lipidomic changes as in the cerebral hemisphere (**Fig. S1**). In addition, the rest of the E10.5 embryos after removal of the brain region were also analyzed, revealing a lipotype similar to the brain region at this time-point (with only 13 out of 1219 (1%) analyzed lipids showing a significant two-fold or larger change in abundance between the E10.5 brain and the remaining tissue, **Fig S2**). This suggests that the neural lipotype acquisition has not yet begun as of E10.5, making it a suitable starting point for profiling the lipidomic changes that concur with neural development.

Taken together, our lipidomic resource provides data on the molar abundance of individual lipid species with annotation of individual fatty acyl chains, thereby providing a comprehensive compendium of over a thousand molecular lipid species. We observed that neural development coincides with a step-wise lipotype acquisition. Specifically, our results reveal that 18:0-sphingolipids and 22:6-glycerophospholipids (i.e., PS, PE and PE O-) already become enriched *in utero*, which coincides with the onset of neurogenesis (Caviness, 1982; Finlay & Darlington, 1995), whereas enrichment of cholesterol occurs postnatally.

### Stem cell-derived neurons do not per se acquire a brain-specific lipotype

The onset of lipotype acquisition during early embryogenesis suggests that this process is coupled to the differentiation events that occur during early brain development. In order to investigate this in a model system that is readily amenable to genetic and pharmacological perturbations, we turned to *in vitro* differentiation of stem cells into neurons. To this end, we made use of an established model where mESCs are grown as suspended cellular aggregates and differentiated into neurons over a time course of 12 days, as confirmed by their neuronal morphology and expression of the neuronal marker β−tubulin III (β-tub III, **Fig. 2A** and **B**) (Bibel et al., 2007). 84 ± 1 % cells were positive for β-tub III, indicating high purity of differentiated neurons on day 12.

**Figure 2.**
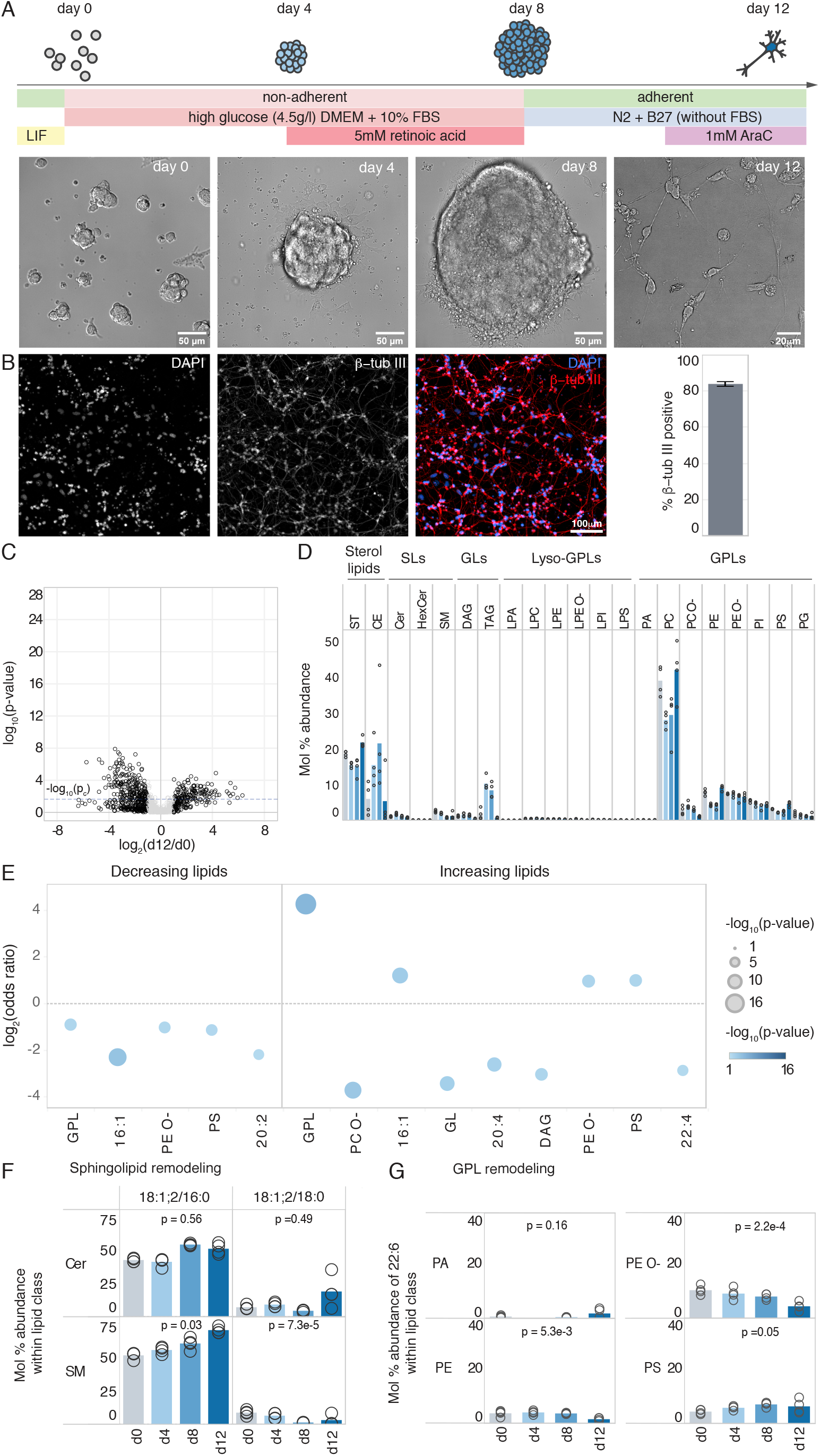
Lack of lipotype acquisition during neuronal differentiation *in vitro*. **A)** Schematic of the neuronal differentiation protocol for mESCs with representative bright-field images at stages in the differentiation protocol where samples were collected for lipidomics. **B)** Representative fluorescence images of cells on day 12 stained for DAPI and the neuronal marker βTub-III are shown. Quantification of βTub-III positive cells is shown on the right, based on 16 images from n = 2 biological replicates. **C)** Volcano plot of molecular lipid species. The fold-change in mol% of each molecular lipid species is calculated between day 0 and day 12 of differentiation. **D)** Profile of different lipid classes across time. **E)** Lipid feature ENrichment Analysis (LENA) of the pool of decreasing and increasing lipids. **F)** Profile of sphingolipids of interest within their respective lipid classes. **G)** Profile of 22:6 (DHA) among fatty acyls constituting PA, PE, PE O- and PS species. Data was collected on n = 4 replicates. p-values were obtained using a one-way ANOVA test and changes were considered significant if p ≤ pc= 0.023 as determined by the Benjamini-Hochberg procedure.

Along this timeline, we analyzed cells by MS^ALL^ lipidomics at 0, 4, 8 and 12 days after the onset of differentiation. Overall, the lipidomic analysis afforded quantitative monitoring of 1355 lipid molecules encompassing 26 lipid classes (**Table S1**). Of these, the levels of 431 lipids (32%) were significantly altered over the course of differentiation, with a two-fold or greater difference between day 0 and day 12 (ANOVA followed by a Benjamini-Hochberg procedure, p-value ≤ p_c_ = 0.023), with 200 (15%) and 231 (17%) increasing and decreasing, respectively (**Fig. 2C**).

At the bulk lipid class-level, we observed a significant increase in the storage lipids CE and TAG on day 4 and day 8, indicating that the cells were storing fat at this stage (**Fig. 2D**). LENA demonstrated that the structural features GPL, 16:1, PE O- and PS were enriched in the pool of increasing lipids (p =8·10^−6^, 1·10^−3^, 1·10^−2^, 1·10^−2^). The increase in 16:1 is indicative of *de novo* lipid synthesis (Lin, 1978; Chakravarty et al., 2004; Freyre et al., 2019), which coincides with the removal of serum from the culture medium and a switch to using glucose as the primary substrate for *de novo* lipogenesis on day 8 (**Table S1**). Further, the polyunsaturated features 20:4 and 22:4 were depleted among the pool of increasing lipids (p = 4·10^−3^, 3·10^−2^) (**Fig. 2E**). Notably, these structural attributes far from resemble the unique lipotype of brain tissue (**Fig. 1E**).

Finally, to explore in further detail this profound discrepancy between lipotype acquisition *in vivo* during neural development and *in vitro* during neuronal differentiation, we inspected the temporal dynamics of the canonical brain lipid biomarkers. This highlighted that only cholesterol is significantly increased, from 19 mol% on day 0 to 22 mol% on day 12 (**Fig. 2D**), albeit not to the same high level as in neural tissue, where the increase is from 21 mol% at E10.5 to 32 mol% at P21. Notably, all 22:6-glycerophospholipids are relatively low in abundance at all time points and a majority of these are also progressively reduced during differentiation (**Fig. 2G** vs. **Fig. 1G**). A similar trend was observed for the sphingolipids Cer 18:1;2/18:0 and SM 18:1;2/18:0, which was offset by an increase in Cer 18:1;2/16:0 and SM 18:1;2/16:0 (**Fig. 2F**). Notably, 16:0-sphingolipids were found to decrease considerably during brain development *in vivo* (**Fig. 1F**). Apart from the lack of lipid remodeling, we also observed that the molecular composition of PIs in the cells differed from that of brain tissue, with elevated levels of 18:1 chains and reduced levels of 20:4 chains in the PI molecules (**Fig. S4**). In summary, our analysis shows that although the differentiated mouse stem cells resemble neurons by morphology and certain protein markers on day 12 of differentiation, they do not *per se* acquire a unique neuronal lipotype akin to brain tissue *in vivo*.

### Lack of canonical lipid markers is a general feature of in vitro neuronal differentiation

Prompted by our finding that neurons generated by culturing mESCs in embryoid bodies (Bibel et al., 2007) do not acquire a brain-like lipotype, we examined lipotype acquisition in two other lineages of stem cell-derived neurons: (1) *in vitro* differentiation of mESCs cultured in an adherent monolayer and stimulated with retinoic acid (Ying et al., 2003) and (2) differentiation of human iPSCs to a mixed population of dopaminergic and GABAergic neurons (Bogetofte et al., 2019). Briefly, the human iPSCs are differentiated to neural stem cells (NSCs) through a neural rosette-based protocol (Swistowski et al., 2009). The NSCs are then further differentiated for up to 25 days to postmitotic neurons with Sonic hedgehog (Shh) stimulation to induce dopaminergic specification (Bogetofte et al., 2019). Neuronal lineage of these cells was confirmed by immunostaining for neuron-specific β-tub III and Microtubule Associated Protein 2 (MAP2) on day 25, where 86.4. ± 1.0 % and 67.9 ± 0.9% of the cells were positive for β-tub III and MAP2, respectively (**Fig. S6A** and **B**).

In both cases, the resulting cells failed to acquire the characteristic lipotype of neural tissue (**Figs. S5** and **S6**). Despite presenting neuronal morphological features, expressing neuron-specific mRNA and protein biomarkers e.g., β-tub III, Map2, Sox1, NeuN), and having synaptic activity (Bibel et al., 2007; Bogetofte et al., 2019). These findings suggest a general failure of *in vitro* neuronal differentiation models in prompting cells to acquire the characteristic lipotype of brain tissue and primary neurons. This discrepancy highlights a potential limitation of using *in vitro* differentiated neurons for mechanistic studies of lipid metabolic programming and lipotype acquisition. Moreover, it prompts the need to develop new *in vitro* differentiation protocols that can adequately commit stem cells to acquire a specific neural lipotype. We note here that the standard culture media for *in vitro* neuronal differentiation of mESCs, containing the B27 supplement, are devoid of the essential polyunsaturated fatty acids DHA (22:6ω3) and arachidonic acid (20:4ω6). Instead, the B27 supplement contains their respective precursors, linolenic acid (18:3ω3) and linoleic acid (18:2ω6) (Brewer & Cotman, 1989; Brewer et al., 1993).

### *Fatty acid supplementation partially establishes a brain-like lipotype* in vitro

Studies using primary rat neurons and astrocytes have previously shown that the polyunsaturated fatty acids 22:6ω3 and 20:4ω6, found to be abundant in brain lipids, are produced by astrocytes from the precursors, 18:3ω3 and 18:2ω6, and thereafter salvaged by neurons (Moore et al., 1991; Kim, 2007). This indicates that the external supply of fatty acids to *in vitro* differentiated stem cells is important for lipid remodeling and accretion of a neuronal lipotype. Thus, we attempted to recapitulate the lipotype acquisition observed *in vivo* by supplementing the culture media of differentiating mESCs with relevant brain lipid precursors. Specifically, we supplemented the cells with a mix of 20 μM DHA (for the production of 22:6-PE, PE O- and PS lipids), 10 μM arachidonic acid (for the biosynthesis of 20:4-PI lipids) and 30 μM stearic acid (for the production of 18:0-sphingolipids). We added the fatty acid mixture to the cell culture media from day 8 of differentiation (**Fig. 3A**). We then collected the fatty acid-supplemented cells as well as untreated control cells on day 12 for lipidomics.

**Figure 3.**
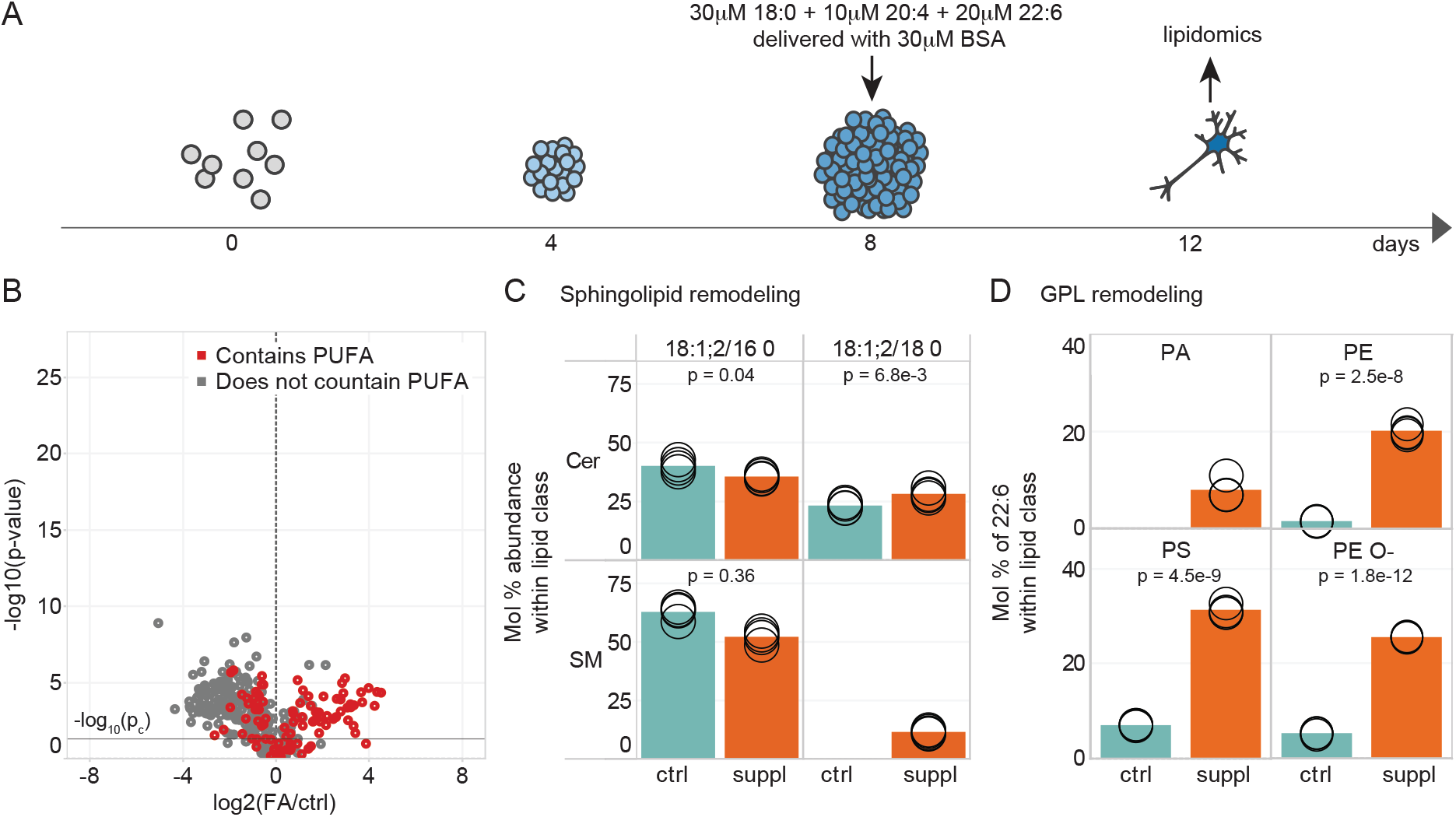
Incorporation of fatty acids into the cell’s lipotype. **A)** Schematic of the differentiation protocol where cells were supplemented with fatty acids. **B)** Volcano plot of molecular lipid species. The fold-change in mol% abundance of each molecular lipid species was calculated between the FA-supplemented and control cells. Red circles correspond to lipid species containing 4 or more double bonds. **C)** Profile of Cer and SM 18:1;2/16:0 and 18:1;2/18:0 in control (cyan bar) and FA-supplemented (orange bar) cells on day 12 of differentiation. **D)** Profile of 22:6 among fatty acyls constituting PA, PE, PE O- and PS species. Data was collected on n = 4 replicates. p-values were obtained using Student’s t-test and considered significant if p ≤ p_c_ = 0.029, as determined by the Benjamini-Hochberg procedure.

The lipidomics analysis identified and quantified 1200 lipid molecules from 26 lipid classes (**Table S1**). We found that among the 310 lipid molecules (26%) significantly altered in abundance by two-fold or more (t-test followed by Benjamini-Hochberg procedure, p-value ≤ p_c_ = 0.029), 124 (10%) were increased and 186 (16%) were reduced. 112 (90%) of the increased lipids likely featured a polyunsaturated chain (using the criteria that at least one of the acyl chains had ≥ 4 double bonds or that the total number of double bonds in the lipid ≥ 4 for lipids whose acyl chain composition could not be determined) (**Fig. 3B**). The incorporation of 22:6 into PE, PE O- and PS closely resembled that seen in the P21 brain, with the molar abundance of 22:6 reaching 20 mol% in PE, 25 mol% in PE O- and 31 mol% in PS (as compared to 22, 22 and 33 mol% in the P21 brain, respectively) (**Fig. 3D**). The incorporation of 20:4 into PI also resembled that seen in brain, with the molar abundance of 20:4 reaching 38 mol% (as compared to 43 mol% in the P21 cerebral hemisphere) (**Fig. S7B)**. The stearic acid supplementation promoted only a modest increase in 18:0-containing sphingolipid species, reaching 28 mol% for ceramides and 12 mol% for SMs (as compared to 76 mol% for both ceramides and SMs in the P21 cerebral hemisphere) (**Fig. 3C**). We note that fatty acid supplementation did not alter the bulk level of individual lipid classes (**Fig. S7A)**.

In summary, it appears that the lipid metabolic machinery underpinning the brain lipotype is partially established in the *in vitro* differentiated neurons. On one hand, it is correctly programmed to be able to take up and incorporate the essential polyunsaturated 22:6 and 20:4 into membrane glycerophospholipids. On the other hand, the metabolic branch responsible for sphingolipid production appears to require more rewiring as it fails to produce the high levels of 18:0-containing sphingolipids seen in developed brain tissue. Together, our results pinpoint two key factors responsible for lipotype acquisition, namely cell-intrinsic enzymatic activities of the underlying lipid metabolic machinery and cell-extrinsic lipid building blocks such as polyunsaturated fatty acids (i.e., derived from the cell culture medium *in vitro* and the neighboring cell types in tissues *in vivo*).

## Discussion

In this study, we investigated the brain lipidome from a developmental perspective, starting at the early embryonic stage. This resource compliments previous lipidomic investigations of brain regions, cell types and membrane fractions obtained from postnatal pups and adolescent animals (Breckenridge et al., 1972; Dawson, 2015; Lauwers et al., 2016; Tulodziecka et al., 2016; Fitzner et al., 2020), and adds to the emerging multi-omics picture of neural development (Yousefi et al., 2021). Our time series analysis allowed us to identify brain lipid biomarkers that increase in abundance throughout development, starting *in utero* (22:6-glycerophsopholipids and 18:0-sphingolipids) while cholesterol increases postnatally (**Fig. 1**). The lipotype acquisition that we observe in the developing mouse brain underscores the important role of lipids and their metabolism in the acquisition of function *in vivo*. To investigate this further, we turned to *in vitro* neuronal differentiation of mESCs and human iPSCs following established protocols (Ying et al., 2003; Bibel et al., 2007). We did not find lipidomic changes supporting the acquisition of neuronal identity, despite the establishment of a neuronal morphology and the expression of neuronal markers (**Figs. 2, S5 and S6**) (Izant & McIntosh, 1980; Haendel et al., 1996; Katsetos et al., 2003; Daubner et al., 2011). Two factors could result in impaired lipotype acquisition in this case, namely inappropriate programming of the underlying lipid metabolic machinery required for the biosynthesis of neural lipids, the absence of required metabolic precursors, or a combination thereof. It has previously been shown that DHA (22:6) is produced in astrocytes and is thereafter salvaged by neurons (Moore et al., 1991; Kim, 2007). Our fatty acid supplementation confirms that the lack of 22:6-glycerophospholipids is indeed due to a deficiency in the culture medium, as supplementation with 20 μM DHA allows cells to produce 22:6-glycerophospholipids to a level comparable with the brain lipotype (**Fig. 3**). On the other hand, supplementing the cells with stearic acid (18:0) does not result in high levels of 18:0-sphingolipids. It is known that the Cer synthase CerS1 is specific for stearoyl-CoA (Venkataraman et al., 2002), which results in the production of 18:0-sphingolipids, while the synthases CerS5 and CerS6 are responsible for 16:0-sphingolipid production. During brain development, one observes a 35-fold increase in the expression of CerS1 and a downregulation of CerS5 and CerS6 compared to embryonic tissue (Sladitschek & Neveu, 2019) (**Fig. S8A)**. In contrast, during *in vitro* neuronal differentiation, between day 8 and 12, CerS1 expression increases only by 5-fold and, contrary to expectation, CerS6 expression is upregulated and CerS5 expression is unchanged (Gehre et al., 2020) (**Fig. S8B**). This could underpin the observation that 16:0-sphingolipids remain elevated whereas brain-specific 18:0-sphingolipids only increase marginally, despite supplementation with stearic acid. Overall, this suggests that appropriate programming of the sphingolipid metabolic machinery is not fully established in stem cell-derived neurons.

Previous work has indicated that homeoviscous adaptation can lead to an increase in cholesterol levels to preserve membrane packing when cells of various cell types, including neurons, are supplemented with high levels of PUFAs (Sinensky, 1974; Ernst et al., 2016; Levental et al., 2020). This finding is of particular interest in the context of the brain and its constituent cell types, as they have relatively high levels of cholesterol as well as PUFAs (**Figs. 1 and S9**) (Tulodziecka et al., 2016; Fitzner et al., 2020). However, our data show that cholesterol increases only postnatally and not concomitantly with the increase in PUFAs seen throughout development. Moreover, fatty acid supplementation did not lead to an increase in cholesterol in the differentiating mESCs (**Fig. S7A**), in line with some other cell types (Zech et al., 2009).

Although the cholesterol level increases in some of the standard protocols used in this study (**Figs. 2C and S5B**), the final levels are far from those observed *in vivo*. Previous work has shown that astrocytes are producers of cholesterol and neurons are recipients (Pfrieger & Ungerer, 2011; Valenza et al., 2015; Ferris et al., 2017), again highlighting the importance of lipid exchange between cell types in the brain.

In summary, here we have outlined the lipidomic landscape of brain development in mice by tracing the abundances of over a thousand lipids over developmental time. This has allowed us to identify two key changes that begin in the embryonic phase of development; namely, an increase in 22:6-glycerophospholipids and the replacement of 16:0-sphingolipids by 18:0-sphingolipids. In contrast, we find that the cholesterol level in the brain increases only postnatally. Such orchestrated lipidome remodeling suggests that the differentiation and maturation of brain cells *in vivo* are coupled to lipid metabolic changes. Our attempt to recapitulate this *in vitro* using neuronal differentiation models to study the molecular mechanisms responsible for lipid metabolic commitment has demonstrated that acquisition of brain lipid hallmarks is *per se* absent in *in vitro* generated neurons. To address these deficits, we added metabolic precursors for producing the canonical brain lipids, but this only partially established a brain-like lipotype. Thus, future work is needed to systematically improve *in vitro* neuronal differentiation protocols, and perhaps consider the need for co-cultures, if they are to be used for mechanistic investigations of lipid biochemistry, membrane biology and biophysics. Our approach provides an important framework for further optimization of differentiation protocols to make *in vitro* neurons more closely recapitulate their *in vivo* counterparts.

## Materials and Methods

### Chemicals and lipid standards

Chloroform, methanol and 2-propanol (Rathburn Chemicals, Walkerburn, Scotland), and ammonium formate (Fluka Analytical, Buchs, Switzerland) were all HPLC grade. Lipid standards were purchased from Avanti Polar Lipids (Alabaster, AL, USA) and Larodan Fine Chemicals (Malmö, Sweden).

### Neuronal differentiation of mESCs in embryoid bodies in vitro

mESCs (129XC57BL/6J generated from male 129-B13 agouti mice) were initially cultured on a feeder layer of mouse fibroblast cells from CD1 mice in ESC media containing Knockout-DMEM (Gibco, Cat. #10829018) with 15% EmbryoMax FBS (Merck, Cat. #ES009-M) and 20 ng/ml leukemia inhibitory factor (LIF from EMBL Protein Expression and Purification Core Facility), 1% non-essential amino acids (Gibco, Cat. #11140050), 1% Glutamax (Gibco, Cat. #35050061), 1mM Sodium pyruvate (Gibco, Cat. #11360070), 1% (50U/ml) Pen/Strep (Gibco, Cat. #15070063) and 143μM β-mercaptoethanol (Gibco, Cat. #21985023). They were cultured over three passages after which the feeder cells were selectively depleted from culture by allowing them to adhere on tissue culture dishes for 10 minutes. The unadhered mESCs were replated for further propagation where the they were differentiated into neuronal progenitor cells over the course of 12 days according to the protocol in Bibel et al., 2007 (**Fig. 2a**). For this, the mESCs were grown in suspension in non-adherent dishes containing differentiation media with high glucose DMEM (Gibco, Cat #11965092), 10% FBS (Gibco, Cat. #26140079) and no LIF, but otherwise identical in composition to the ES medium. This results in the formation of embryoid bodies that grow larger over the course of 8 days. On day 4, 5 μM retinoic acid was added to induce neuronal differentiation. On day 8, Embryoid bodies were dissociated with trypsin (Gibco, Cat. #25300054) and cells were plated at a density of 140,000 cells/cm^2^ on plates pre-coated with poly-D-lysine (PDL) and Laminin 511 (Biolamina, Cat. #LN511) in N2B27 media (High glucose DMEM supplemented with 1% N2 (Gibco, Cat. #17502048), 1% B27 (without vitamin A, Gibco, Cat. #17504044), 1% (50U/ml) Pen/Strep and 1 mM Sodium pyruvate (Gibco, Cat. #11360070)). On day 10, cells were treated with 1 μM cytosine arabinoside (AraC, Merck, Cat. #C1768) to kill proliferating, non-differentiated cells.

Media was changed every day before day 0 due to rapid cell proliferation in the stem cell state and every two days thereafter, with special care to prevent dissociation of the embryoid bodies due to shear and care to avoid exposing the plated neuronal progenitors to air. Cells were placed at 37°C with 5% CO_2_ under all media conditions.

### Fatty acid supplementation

Stock solutions of fatty acids to be supplemented were prepared in ethanol (40 mM 18:0, 32.8 mM 20:4 and 60 mM 22:6) and pipetted dropwise into N2 media containing 3 mM fatty acid-free BSA (Merck, Cat. #A8806) at 37°C with constant stirring to obtain final concentrations of 3 mM 18:0, 1 mM 20:4 and 2 mM 22:6. This 100x stock of the FA-supplemented N2 media was aliquoted and stored at -20°C. On day 8 of the differentiation protocol, it was added to the N2 culture medium in a 1:100 ratio.

### Neuronal differentiation of mESCs in a monoloyer in vitro

mESCs were maintained as undifferentiated stem cells in an adherent monolayer in N2B27 media (a 1:1 mixture of DMEM/F12 (without HEPES and with glutamine, Gibco, Cat. #131331028) and Neurobasal medium (Gibco, Cat. #21103049) supplemented with 0.5x N2 (Gibco, 17502001), 0.5x B27 (without vitamin A), 0.25mM L-glutamine (Gibco, Cat. #25030149), 0.1 mM β-mercaptoethanol (Gibco, Cat. #21985023), 10 mg/ml BSA fraction V (Merck, Cat. # 10735078001), 10 mg/mL Human recombinant Insulin (Merck, Cat. #91077C) and 1% Pen/Strep) +2iLIF (10 ng/ml leukemia inhibitory factor + 3 μM CHIR99021 + 1 μM PD0325901 from Tocris, Cat. # 4423 and 4192, respectively). The cells were grown on 0.1% gelatin-coated dishes and usually seeded at a density of 0.5–1.5 × 10^4^/cm^2^. Medium was replaced every day and the cells were trypsinized and reseeded every two days in their undifferentiated state. For monolayer differentiation, the cells were similarly seeded and kept in N2B27 without the 2iLIF for 24 hours, following which the media was supplemented with 1 μM retinoic acid and refreshed every 24 hours.

### Neuronal differentiation of human iPSCs in vitro

The human iPSC line XCL-1 (XCell Science Inc., Novato, CA, USA) was differentiated to NSCs by XCell Science Inc. using a 14-day protocol where iPSCs were initially differentiated in suspension as embryoid bodies followed by the formation of attached neural rosettes. These NSCs were then isolated and expanded (Swistowski et al., 2009). The NSCs were propagated using Geltrex (Thermo, Cat. #A1413202) coated plates in Neurobasal Medium (Thermo, Cat. #21103049) supplemented with NEAA (Thermo, Cat. #11140050), GlutaMax-I (Thermo, Cat. #35050038), B27 supplement (Thermo, Cat. #17504044), penicillin-streptomycin (Thermo, Cat. #15140), and basic fibroblast growth factor (bFGF, R&D Systems, Cat. #233-FB). Cells were enzymatically passaged with Accutase (Thermo, Cat. #A1110501) when 80-90% confluent. Neuronal differentiation was achieved by culturing NSCs in DOPA Induction and Maturation Medium (XCell Science, Cat. #D1-011) according to manufacturer instructions by supplementing with 200 ng/ml human recombinant Sonic Hedgehog (Peprotech, Cat. #100-45) from day 0-10 and passaging cells at day 0, 5 and 10 onto poly-L-ornithine (Sigma, Cat. #3655) and laminin (Thermo, Cat. #23017015) coated plates at a density of 50,000 cells/cm^2^.

### Immunostaining

mESCs differentiated into neuronal progenitor cells for twelve days were fixed in 4% (v/v) paraformaldehyde (Thermo Fisher, Cat. #28908) for twenty minutes. Excess paraformaldehyde was quenched with 30 mM glycine for five minutes, and coverslips were washed three times with PBS. Cells were permeabilized with 0.1% Triton-X 100 (Carl Roth Cat. # 3051.4) and blocked with 0.5% BSA (Carl Roth Cat. # 8076.4) for thirty minutes at room temperature. Cells were incubated with a primary antibody against β-tubulin III (Abcam Cat. # ab78078) diluted in 0.5% BSA 1:200 and incubated for one hour at room temperature. A secondary antibody conjugated to an Alexa® Fluor 594 dye (Invitrogen, Cat. # A-11005) was used for detection. Cells were stained with DAPI 5 μg/ml (Sigma Cat. # D9542) for five minutes, and coverslips were mounted on glass slides with ProLong Gold (Invitrogen Cat. # P36934). Images were acquired with a Nikon Eclipse Ti fluorescence microscope using the 20x objective.

Human iPSC-derived neurons, differentiation day 25 from NSCs, were fixed for 15 min at RT in 4% (w/v) paraformaldehyde (Sigma, Cat. #158127) and permeabilized with 0.1% Triton-X-100 (Sigma, Cat. #9002-93-1). Unspecific binding was blocked with 10% goat serum (Millipore, Cat. #S26) and cultures were incubated overnight at 4°C with primary antibodies diluted in TBS/10% goat serum: mouse anti-β-tubulin-III (Sigma #T8660) or mouse anti-MAP2 (Sigma, Cat.# M1406). Incubation with secondary antibodies: Alexa Fluor 555 goat anti-mouse IgG (Molecular Probes, Cat. #A21422) were done at 1:500 in TBS/10% goat serum for 2 hr at RT. Cell nuclei were counterstained with 10μM 4”,6-diamidino-2-phenylindole dihydrochloride (DAPI, Sigma, Cat. #D9542). Coverslips were mounted onto glass slides with ProLong Diamond mounting medium (Molecular Probes, Cat. #P3690).

To estimate the proportion of differentiated neurons, the ratio of neuronal marker-positive cells to the total number of cells (DAPI) in each field of view was determined and averaged over several images.

### Sample collection for lipidomics

Brain tissue samples for lipidomics were collected from wild-type CD-1 mice from the EMBL breeding colonies at four different time points in development, namely the embryonic stages E10.5 and E15.5 and the post-natal stages P2 and P21 (**Fig. 1A**). No procedure was performed on live animals. The mothers carrying E10.5 and E15.5 embryos, and P2 and P21 pups were euthanized by IACUC approved methods, after which tissues were collected for lipidomics.

At E10.5, each brain sample consisted of pooled brain region from five embryos in order to obtain enough sample for lipidomics. At all remaining time points, the brain from a single mouse embryo or pup was dissected to obtain two samples: one with cerebral hemispheres and another with the rest of the brain. All tissues were washed with PBS, flash-frozen in liquid nitrogen and stored at -80°C till lipid extraction.

Mouse cells were collected for lipidomics on days 0, 4, 8 and 12 of differentiation in embryoid bodies of mESCs. On day 0, cells were collected following trypsinization. On days 4, 8 and 12, suspended embryoid bodies from one 6 cm dish each were collected by a cell scraper. In the case of monolayer differentiation of mESCs, cells in one well of a six-well plate were collected per sample. In each case, the cells were pelleted, washed twice in HEPES-KOH buffer, suspended in 200-300 μL 155 mM ammonium formate, flash-frozen in liquid nitrogen and stored at -80°C till lipid extraction.

Human iPSC-derived neurons were collected following treatment with Accutase on differentiation days 0, 10 and 25 (from the NSC stage). Cells were collected in ice cold PBS, pelleted, washed with 155 mM ammonium acetate (Sigma, Cat. #A7330), flash-frozen in liquid nitrogen and stored at -80°C till lipid extraction.

### Lipid extraction

Lipids were extracted from the samples as described previously (Almeida et al., 2014). In short, all samples were thawed at 4°C and homogenized by sonication (and mechanically using an ultra-turrax in addition to this, in the case of brain samples). Samples were spiked with an internal standard (IS) mix prepared in-house and containing known amounts of synthetic lipid standard (**Table S1**). Lipid extraction was performed by 2-step extraction (Sampaio et al., 2011), first by partitioning the sample between 155 mM aqueous ammonium formate and chloroform/methanol (10:1, vol/vol) and then using the aqueous fraction to partition against chloroform/methanol (2:1, vol/vol). The solutions were shaken at 1400 rpm at 4°C during both steps (for 2 hours and 1.5 hours, respectively) and their organic phases collected. Solvents were then removed by vacuum evaporation, leaving deposits of lipids extracted in the 10:1- and 2:1-extracts, respectively. The 10:1- and 2:1-extracts were dissolved in chloroform/methanol (1:2, vol/vol).

### Shotgun lipidomics

MS^ALL^ lipidomics analysis was performed as previously described (Almeida et al., 2014). In short, mass spectra of the lipid extracts were recorded in both positive and negative ion modes using an Orbitrap Fusion Tribrid mass spectrometer (Thermo Fisher Scientific) equipped with a robotic nanoflow ion source, TriVersa NanoMate (Advion Biosciences). Aliquots of 10:1-extracts were diluted with 2-propanol to yield an infusate composed of chloroform/methanol/2-propanol (1:2:4, vol/vol/vol) and 7.5 mM ammonium formate for positive ion mode analysis. The 10:1-extracts were also diluted with 2-propanol to yield an infusate composed of chloroform/methanol/2-propanol (1:2:4, vol/vol/vol) and 0.75 mM ammonium formate for negative ion mode analysis. Aliquots of the 2:1-extracts were diluted with methanol to yield a infusate composed of chloroform/methanol (1:5, vol/vol) and 0.005% methylamine for analysis in negative ion mode. High-resolution mass spectra of intact ions (MS1) were recorded using a the orbitrap mass analyzer and precursor ions from each 1 Da window within the detected range were fragmented and the mass spectra of the fragments (MS2) were recorded using the orbitrap mass analyzer. Lipid molecules and fragment ions were identified using ALEX^123^ (Pauling et al., 2017; Ellis et al., 2018). Molar abundance of lipid molecules were quantified by normalizing their MS1 intensities to that of spiked-in internal lipid standards and scaling by the known concentration of the internal standard (**Table S1**). Lipid quantification and downstream analysis of fatty acid composition and Lipid feature ENrichment Analysis (LENA) were carried out on the SAS9.1 platform (SAS) as described previously (Sprenger et al., 2021).

### Statistical analysis

In the analysis of time-course data, replicates from identical conditions were considered to constitute a sample group (resulting in nine sample groups among the mouse brain samples and four sample groups among cell culture samples). Analysis of variance (ANOVA) testing with these sample groups was performed to identify significant changes in the mol% abundances of lipids. An unpaired Student’s t-test was used to test for significant differences in the lipidome of day 12 neurons between control and fatty acid supplemented conditions.

To minimize false positives due to multiple hypothesis testing, we calculated a Benjamini-Hochberg critical value q for every p-value with false positive rate of 0.05 and consider changes significant for all p ≤ p_c_, where p_c_ is the largest p-value that is smaller than its corresponding q-value. The statistical tests were performed using R and the Principal Component Analysis was done on ClustVis (Metsalu & Vilo, 2015). Tableau Desktop (Tableau Software) was used for data visualization.

## Acknowledgements

We thank Martin Hermansson for useful discussions on lipidomics, Emilia Esposito, Juan Carlos Boffi and Vikram Singh Ratnu for providing us with dissected mouse brain tissues, and Eva Pillai and Martin Bergert for critical reading of the manuscript. This research was supported by the Danish Council for Independent Research | Natural Sciences (DFF - 6108- 00493) and the VILLUM Center for Bioanalytical Sciences (VKR023179) to C.S.E., and the European Molecular Biology Laboratory (EMBL) to K-M. N., C.S.E. and A.D-M. H.B. and M.M. were supported by the Innovation Fund Denmark (BrainStem), the Danish Parkinson Foundation, the Jascha Foundation, the A.P. Møller Foundation for the Advancement of Medical Science, and the Faculty of Health Sciences at the University of Southern Denmark.

## Supplementary figures

**Figure S1.**
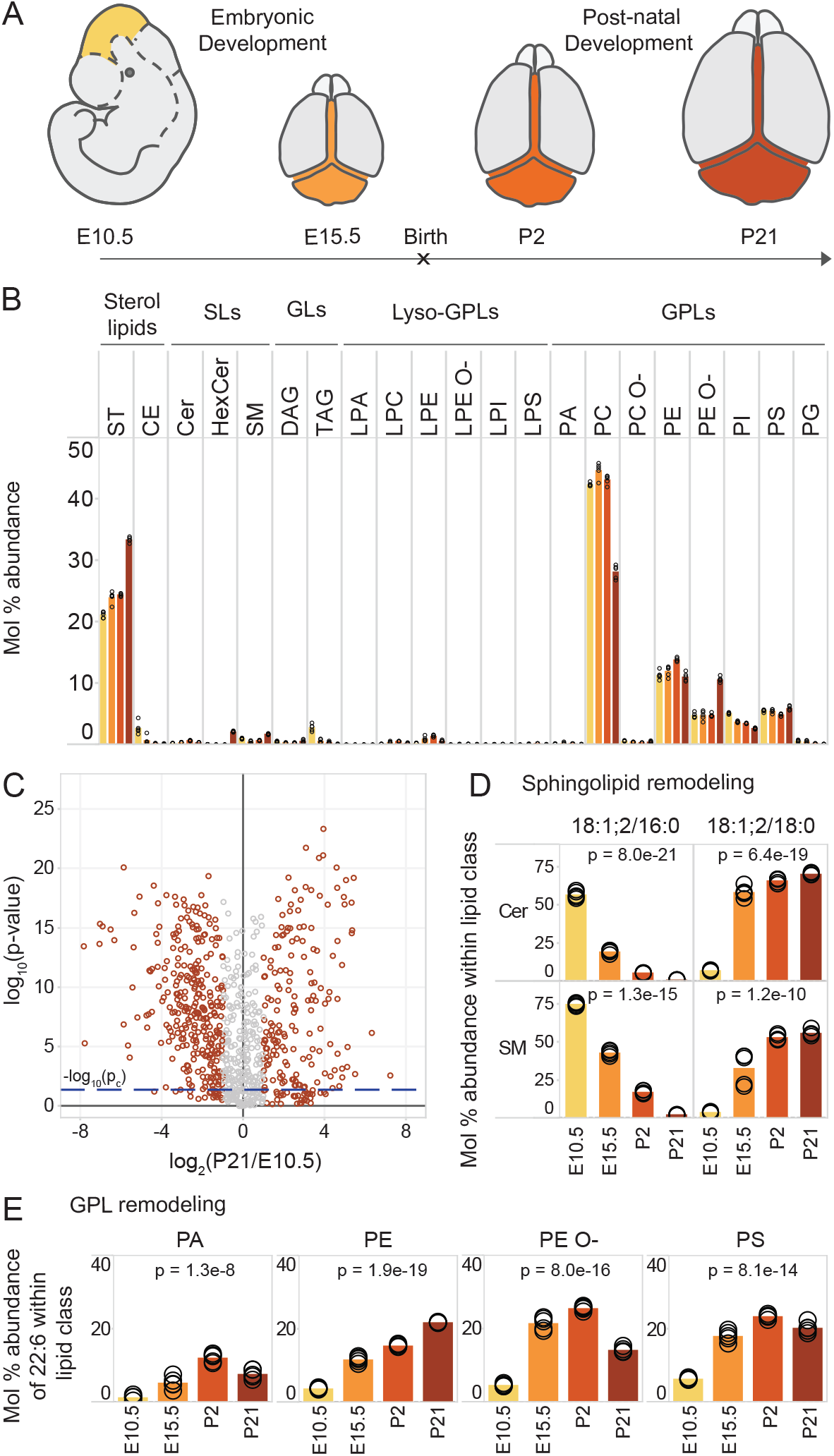
Lipotype acquisition in the developing mouse brain. **A)** Brain regions apart from cerebral hemispheres were collected separately at the indicated time points and analyzed by in-depth MS^ALL^ lipidomics. B) Abundance of different lipid classes across developmental time. Bars representing the mean value for each sample group are colored in accordance with panel A. **C)** Volcano plot of molecular lipid species. The fold-change in mol% of each lipid within its lipid class is calculated between the E10.5 brain region and the rest of the P21 brain after removal of the cerebral hemispheres. **D)** The most abundant ceramide (Cer) and sphingomyelin (SM) species across samples, containing an 18:1;2 sphingoid chain and either a 16:0 or 18:0 acyl chain. **E)** Abundance of 22:6 (DHA) among fatty acyls constituting phosphatidic acid (PA), phosphatidylethanolamine (PE and PE O-) and phosphatidylserine (PS). Data was collected on n = 5 replicates. p-values were obtained using a one-way ANOVA test and considered significant if p ≤ p_c_ = 0.041, as determined by the Benjamini-Hochberg procedure.

**Figure S2.**
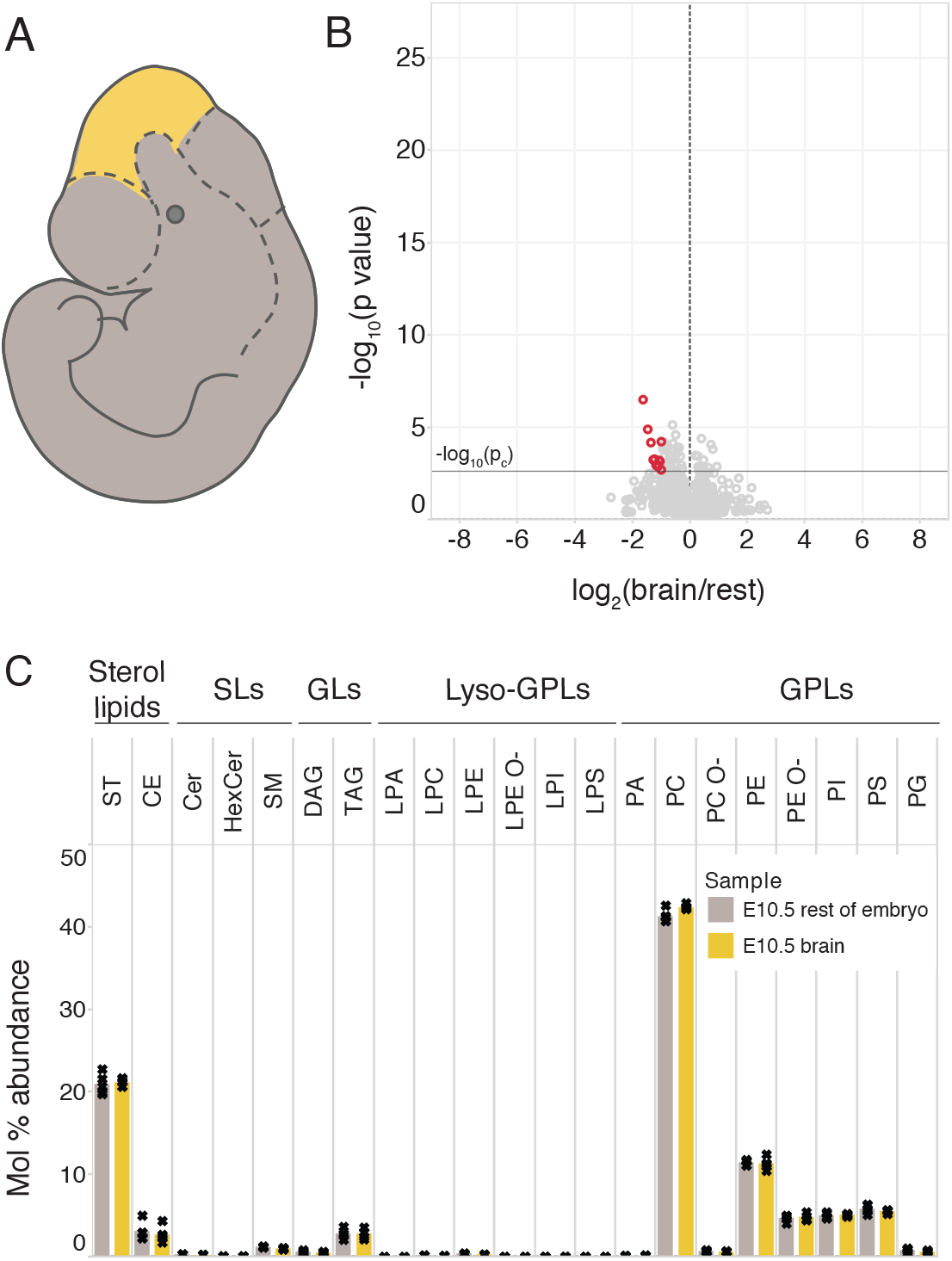
The E10.5 brain lipidome does not significantly differ from that of the rest of the E10.5 embryo. The rest of the E10.5 embryo after excision of the brain region was collected and analyzed by lipidomics. **A)** Schematic showing the dissection and resulting portions of the E10.5 embryo in the two sample groups. **B)** Volcano plot where each circle is one lipid species and its abundance is compared between the E10.5 brain region and the rest of the E10.5 embryo. Lipids with p ≤ p_c_ = 0.0025 (Student’s t-test followed by the Benjamini-Hochberg procedure) and fold-changes greater than two-fold are colored red. **C)** Lipid class profile of the two tissue samples collected at E10.5. Data was collected on n = 5 replicates.

**Figure S3.**
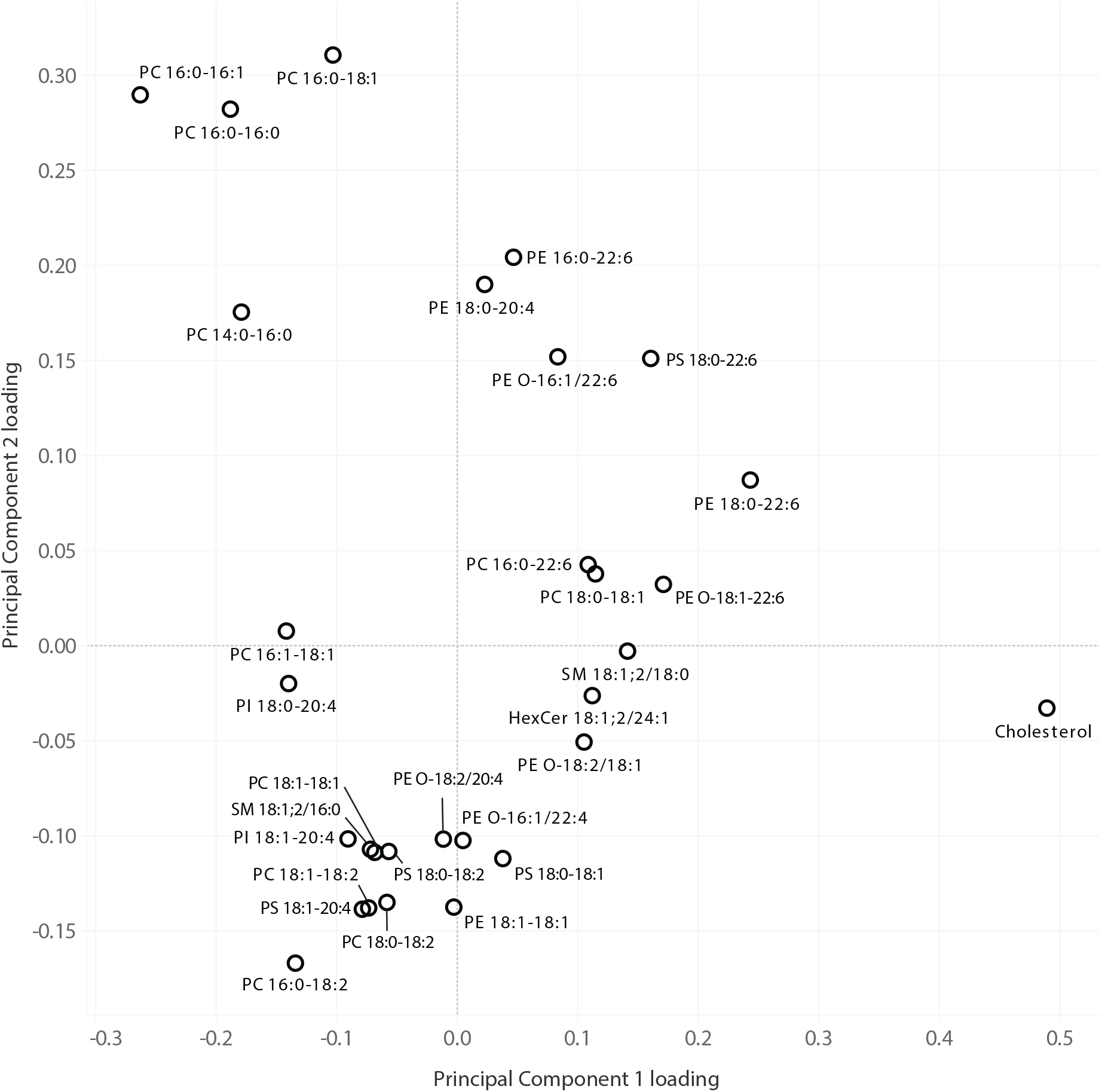
Principal component loadings of lipids in the analysis of mouse tissues. PC1 and PC2 loadings of lipids where either loading was greater than 0.1 in magnitude are shown in this plot.

**Figure S4.**
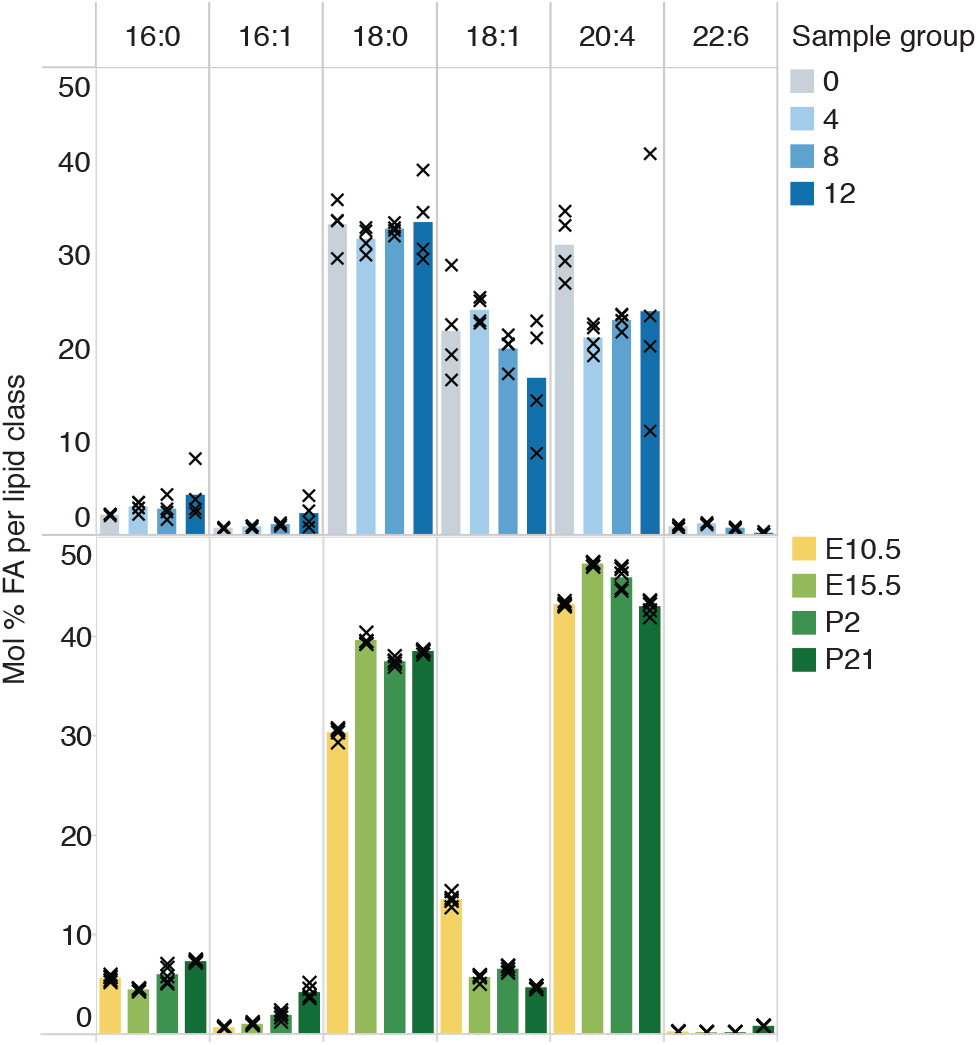
Fatty acid profile of PI lipids. Comparison of the mol% abundances of various fatty acyl chains among PIs in mESCs being differentiated *in vitro* into neurons and mouse brain tissues over development. Data was collected on n = 4 replicates of cells and n = 5 replicates of brain tissue.

**Figure S5.**
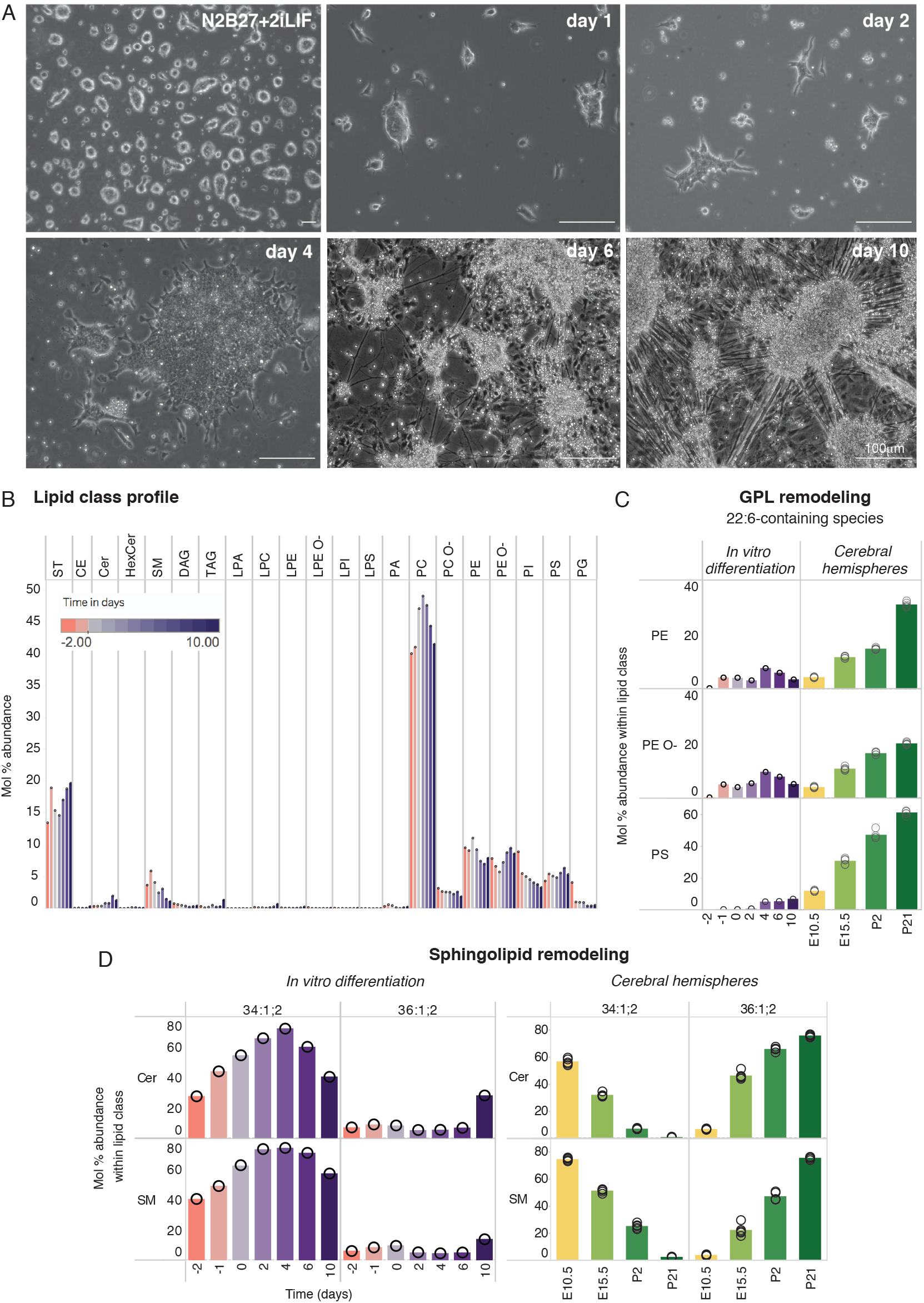
Lipotype acquisition in neuronal differentiation of adherent mESCs. **A)** Neuronal differentiation of adherent mESCs. **B)** The relative abundance of different lipid classes across time. Samples from day -2, -1 and 0 are all in N2B27+2iLIF and are collected 2, 1, and 0 days before the removal of 2iLIF and start of differentiation. B) The relative abundance of 22:6-glycerophospholipids PE 40:6, PE O-40:7 and PS 40:6; 22:6-containing PA was not detected in this dataset and is therefore not shown. **D)** The mol percent abundances of sphingolipids of interest within their respective lipid classes are plotted across samples. As the data is from a single replicate of the differentiation protocol, results were not tested for significance and are meant to be indicative rather than conclusive. Mol% abundance of 22:6-glycerophospholipids and 18:0-sphingolipids in cerebral hemispheres across development are shown on the right panel in C and D for comparison.

**Figure S6.**
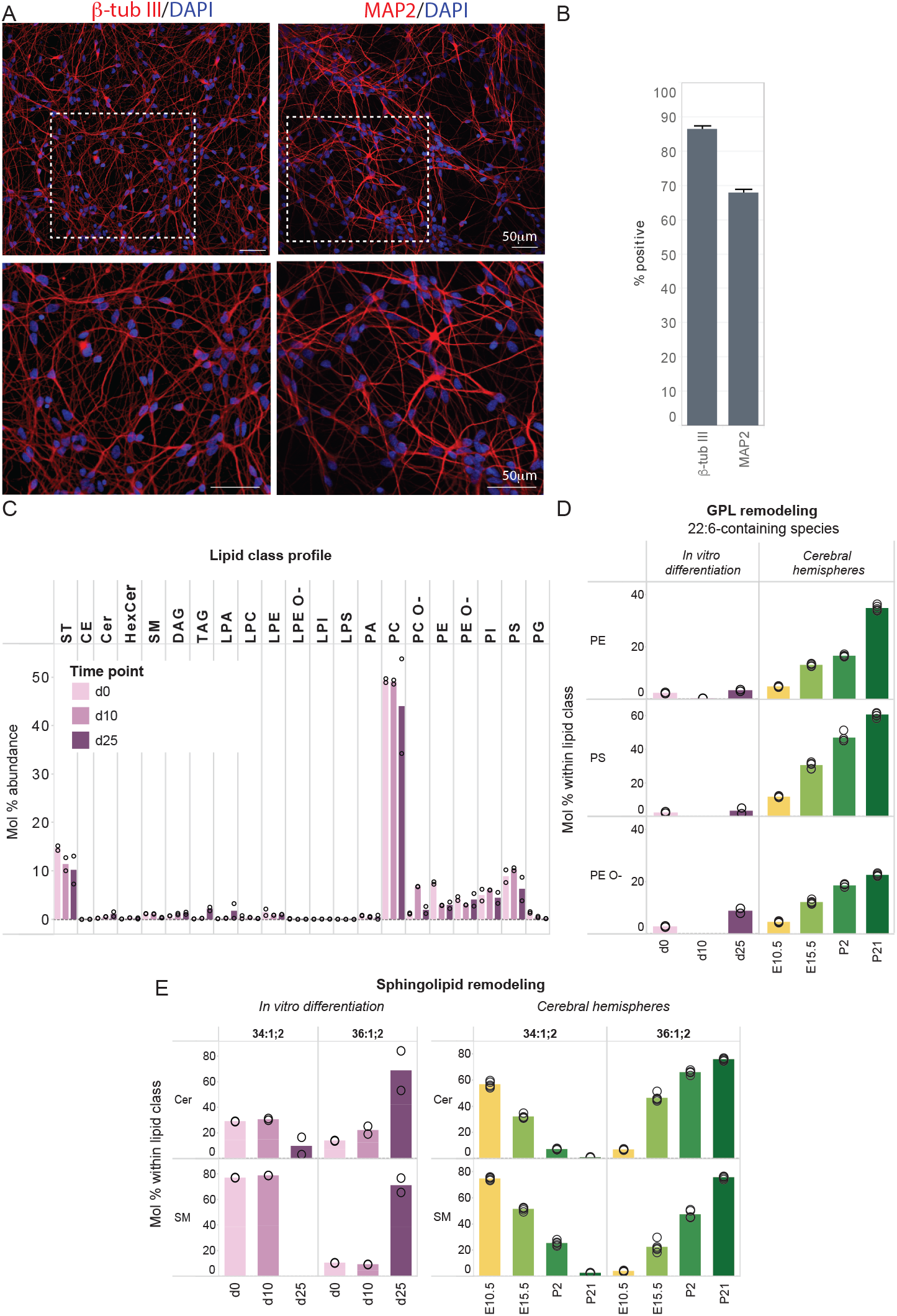
Neuronal differentiation of human iPSC-derived NSCs prompts the acquisition of 18:0-sphingolipids but not 22:6-glycerophospholipids. **A)** Immunofluorescence images of human iPSC-derived neurons on day 25 of differentiation from the NSC stage, labelled for the neuronal markers β-tub III and MAP2. Regions in dashed white lines are shown below. **B)** Percentage of cells positive for β-tub III and MAP2 as quantified from 45 images per group, from n = 3 biological replicates. **C**) Abundance of lipid classes across time from day 0, 10 and 25 of differentiation. **D)** The abundance of the 22:6-glycerophospholipids PE 18:0-22:6, PS 18:0-22:6 and PE O-18:1/22:6; 22:6-containing PA was not detected in this dataset and is therefore not shown. **E)** Abundances of 16:0- and 18:0-sphingolipids; The most abundant Cer and SM species across samples contain a 16:0 fatty acyl chain on day 0 and 10 but are replaced by 18:0-containing Cer and SM, respectively, on day 25. Data was obtained from n=2 independent differentiations. Mol% abundance of 22:6-glycerophospholipids and 18:0-sphingolipids in cerebral hemispheres across development are shown on the right panel in C and D for comparison.

**Figure S7.**
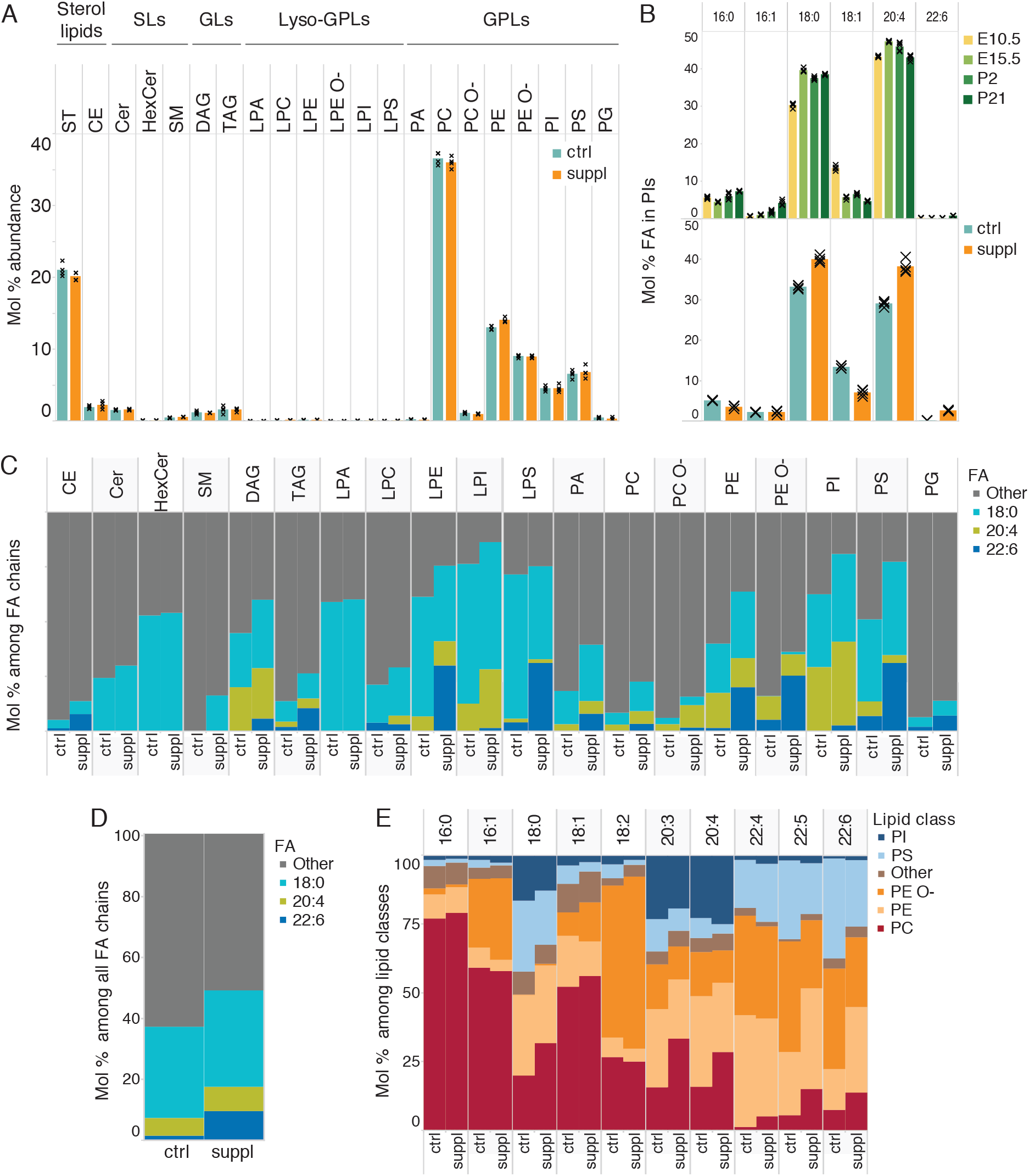
Fatty acid distribution within the cell lipidome. **A)** Profile of lipid classes in control vs FA-supplemented *in vitro* differentiated neurons. **B)** Mol % abundances of various fatty acyl chains among PIs in the developing mouse brain and FA-supplemented *in vitro* differentiated neurons. **C)** Abundance of the three supplemented fatty acids among all those present in individual lipid classes in FA-supplemented cells, showing specificity in the incorporation of the supplemented fatty acids into lipids. **D)** The total abundance of the three supplemented fatty acids in the FA pool of FA-supplemented cells. E) Proportion of abundant fatty acyl chains among the prominent lipid classes in which they are present. Data was collected on n = 4 replicates.

**Figure S8.**
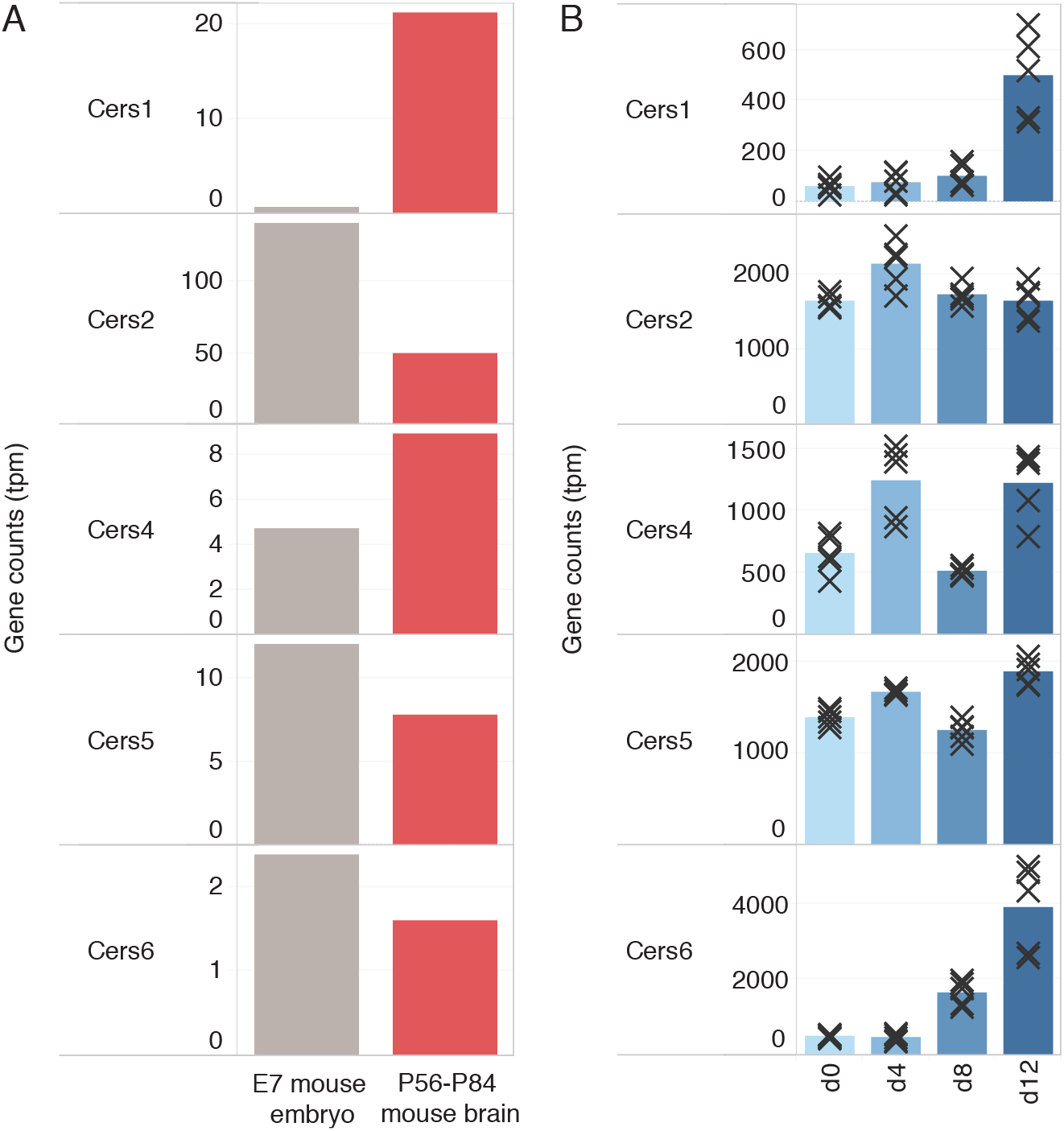
Expression of ceramide synthases involved in sphingolipid biosynthesis. **A)** Expression levels are compared between E7 mouse embryos and pooled brain samples from P56-P84 mice. The data is from n =1 replicate and taken from RNA-seq (Sladitschek & Neveu, 2019). **B)** Expressioon levels are compared across the *in vitro* differentiation of mESCs into neurons in embryoid bodies. The data is from n = 5 replicates and taken from RNA-seq (Gehre et al., 2020). Cers1 and Cers4 are responsible for incorporating C18 chains into sphingolipids whereas Cers5 and Cers6 incorporate C16 chains (Levy & Futerman, 2010).

**Figure S9.**
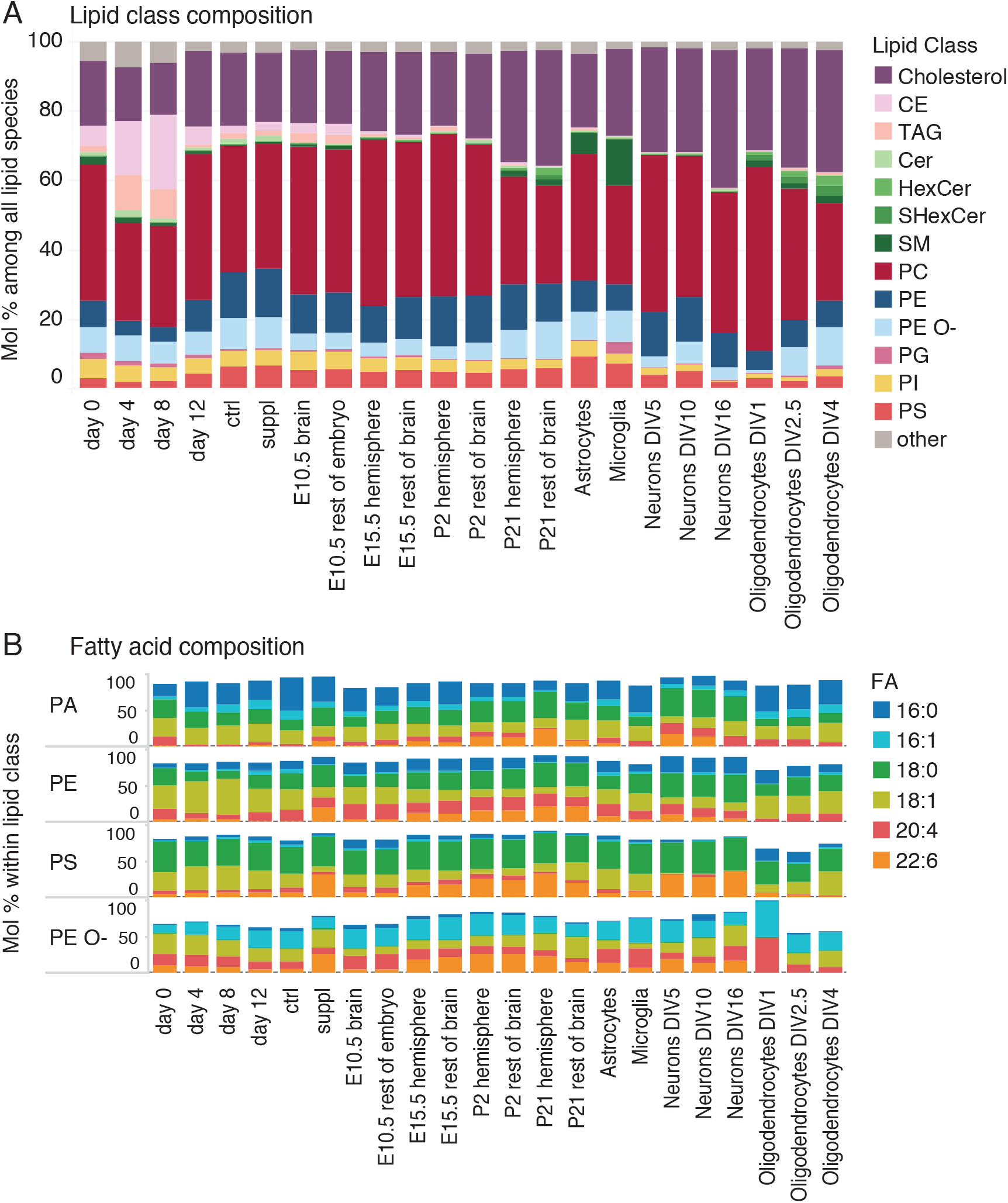
Comparison of lipidomes to primary cells from Fitzner et al, 2020. **A)** Lipid class- and **B)** FA-compositions in all samples compared to primary cells from Fitzner et al, 2020. Samples from *in vitro* differentiation of mESCs were obtained on day 0, 4, 8 and 12 (control and FA-supplemented) of neuronal differentiation in embryoid bodies (Bibel et al., 2007).

## Supplementary table

**Table S1.**
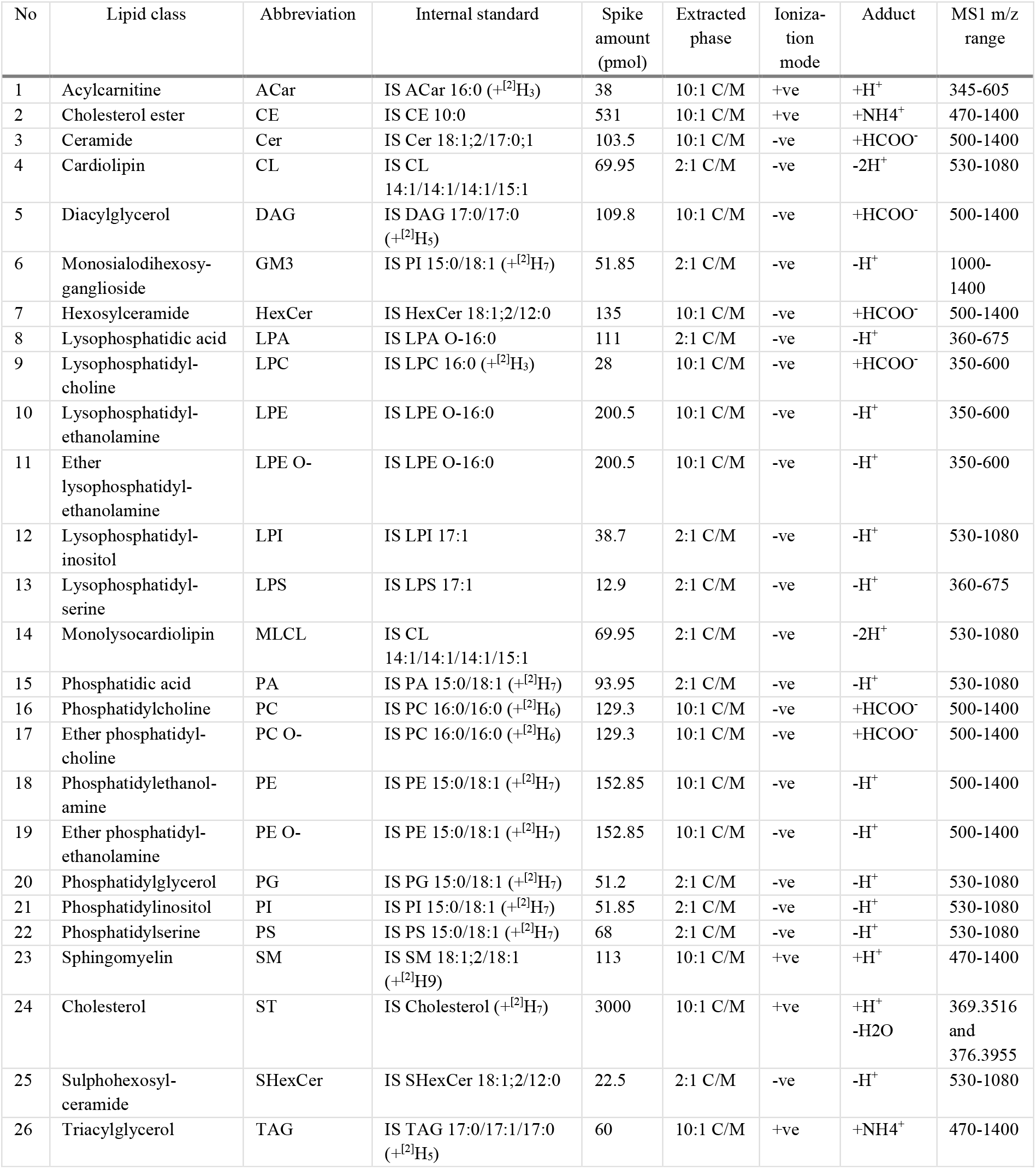
Identification and quantification of lipids of various classes

## Notes

**Conflict of interest statement:** The authors declare that they have no conflict of interest.

### Competing Interest Statement

The authors have declared no competing interest.

### Summary of Updates

The manuscript has been revised as per the suggestions of reviewers in the pre-print review process of Review Commons. Reviewer comments and our response can be found under the TRiP tab.

## References

Almeida, R., Pauling, J. K., Sokol, E., Hannibal-Bach, H. K., & Ejsing, C. S. (2014). Comprehensive lipidome analysis by shotgun lipidomics on a hybrid quadrupole-orbitrap-linear ion trap mass spectrometer. Journal of the American Society for Mass Spectrometry, 26(1), 133–148. https://doi.org/10.1007/s13361-014-1013-x

Bejaoui, K., Wu, C., Scheffler, M. D., Haan, G., Ashby, P., Wu, L., De Jong, P., & Brown, R. H. (2001). SPTLC1 is mutated in hereditary sensory neuropathy, type 1. Nature Genetics, 27(3), 261–262. https://doi.org/10.1038/85817

Bibel, M., Richter, J., Lacroix, E., & Barde, Y. A. (2007). Generation of a defined and uniform population of CNS progenitors and neurons from mouse embryonic stem cells. Nature Protocols, 2(5), 1034–1043. https://doi.org/10.1038/nprot.2007.147

Bieberich, E. (2013). It’s a lipids world. Neurochemichal Research, 37(6), 1208–1229. https://doi.org/10.1007/s11064-011-0698-5

Bogetofte, H., Jensen, P., Okarmus, J., Schmidt, S. I., Agger, M., Ryding, M., Nørregaard, P., Fenger, C., Zeng, X., Graakjær, J., Ryan, B. J., Wade-Martins, R., Larsen, M. R., & Meyer, M. (2019). Perturbations in RhoA signalling cause altered migration and impaired neuritogenesis in human iPSC-derived neural cells with PARK2 mutation. Neurobiology of Disease, 132(August), 104581. https://doi.org/10.1016/j.nbd.2019.104581

Bozek, K., Wei, Y., Yan, Z., Liu, X., Xiong, J., Sugimoto, M., Tomita, M., Pääbo, S., Sherwood, C. C., Hof, P. R., Ely, J. J., Li, Y., Steinhauser, D., Willmitzer, L., Giavalisco, P., & Khaitovich, P. (2015). Organization and Evolution of Brain Lipidome Revealed by Large-Scale Analysis of Human, Chimpanzee, Macaque, and Mouse Tissues. Neuron, 85(4), 695–702. https://doi.org/10.1016/j.neuron.2015.01.003

Breckenridge, W. C., Gombos, G., & Morgan, I. G. (1972). The lipid composition of adult rat brain synaptosomal plasma membranes. BBA - Biomembranes, 266(3), 695–707. https://doi.org/10.1016/0005-2736(72)90365-3

Brewer, G. J., & Cotman, C. W. (1989). Survival and growth of hippocampal neurons in defined medium at low density: advantages of a sandwich culture technique or low oxygen. Brain Research, 494(1), 65–74. https://doi.org/10.1016/0006-8993(89)90144-3

Brewer, G. J., Torricelli, J. R., Evege, E. K., & Price, P. J. (1993). Optimized survival of hippocampal neurons in B27-supplemented neurobasal™, a new serum-free medium combination. Journal of Neuroscience Research, 35(5), 567–576. https://doi.org/10.1002/jnr.490350513

Cao, D., Kevala, K., Kim, J., Moon, H. S., Jun, S. B., Lovinger, D., & Kim, H. Y. (2009). Docosahexaenoic acid promotes hippocampal neuronal development and synaptic function. Journal of Neurochemistry, 111(2), 510–521. https://doi.org/10.1111/j.1471-4159.2009.06335.x

Capolupo, L., Khven, I., Lederer, A. R., Mazzeo, L., Glousker, G., Ho, S., Russo, F., Montoya, J. P., Bhandari, D. R., Bowman, A. P., Ellis, S. R., Guiet, R., Burri, O., Detzner, J., Muthing, J., Homicsko, K., Kuonen, F., Gilliet, M., Spengler, B., … D’Angelo, G. (2022). Sphingolipids control dermal fibroblast heterogeneity. Science, 376(6590). https://doi.org/10.1126/science.abh1623

Caviness, V. S. (1982). Neocortical histogenesis in normal and reeler mice: A developmental study based upon [3H]thymidine autoradiography. Developmental Brain Research, 4(3), 293–302. https://doi.org/10.1016/0165-3806(82)90141-9

Chakravarty, B., Gu, Z., Chirala, S. S., Wakil, S. J., & Quiocho, F. A. (2004). Human fatty acid synthase: Structure and substrate selectivity of the thioesterase domain. Proceedings of the National Academy of Sciences of the United States of America, 101(44), 15567–15572. https://doi.org/10.1073/pnas.0406901101

Daubner, S. C., Le, T., & Wang, S. (2011). Tyrosine hydroxylase and regulation of dopamine synthesis. Archives of Biochemistry and Biophysics, 508(1), 1–12. https://doi.org/10.1016/j.abb.2010.12.017

Davletov, B., & Montecucco, C. (2010). Lipid function at synapses. Current Opinion in Neurobiology, 20(5), 543–549. https://doi.org/10.1016/j.conb.2010.06.008

Dawkins, J. L., Hulme, D. J., Brahmbhatt, S. B., Auer-Grumbach, M., & Nicholson, G. A. (2001). Mutations in SPTLC1, encoding serine palmitoyltransferase, long chain base subunit-1, cause hereditary sensory neuropathy type I. Nature Genetics, 27(3), 309–312. https://doi.org/10.1038/85879

Dawson, G. (2015). Measuring Brain Lipids. Biochimica et Bioiphysica Acta, 1851(8), 1026–1039. https://doi.org/10.1016/j.bbalip.2015.02.007

Ellis, S. R., Paine, M. R. L., Eijkel, G. B., Pauling, J. K., Husen, P., Jervelund, M. W., Hermansson, M., Ejsing, C. S., & Heeren, R. M. A. (2018). Automated, parallel mass spectrometry imaging and structural identification of lipids. Nature Methods, 15(7), 515–518. https://doi.org/10.1038/s41592-018-0010-6

Ernst, R., Ejsing, C. S., & Antonny, B. (2016). Homeoviscous Adaptation and the Regulation of Membrane Lipids. Journal of Molecular Biology, 428(24), 4776–4791. https://doi.org/10.1016/j.jmb.2016.08.013

Ferris, H. A., Perry, R. J., Moreira, G. V., Shulman, G. I., Horton, J. D., & Kahn, C. R. (2017). Loss of astrocyte cholesterol synthesis disrupts neuronal function and alters whole-body metabolism. Proceedings of the National Academy of Sciences of the United States of America, 114(5), 1189–1194. https://doi.org/10.1073/pnas.1620506114

Fewou, S. N., Büssow, H., Schaeren-Wiemers, N., Vanier, M. T., Macklin, W. B., Gieselmann, V., & Eckhardt, M. (2005). Reversal of non-hydroxy: α-hydroxy galactosylceramide ratio and unstable myelin in transgenic mice overexpressing UDP-galactose: Ceramide galactosyltransferase. Journal of Neurochemistry, 94(2), 469–481. https://doi.org/10.1111/j.1471-4159.2005.03221.x

Finlay, B. L., & Darlington, R. B. (1995). Linked regularities in the development and evolution of mammalian brains. Science, 268(5217), 1578–1584. https://doi.org/10.1126/science.7777856

Fitzner, D., Bader, J. M., Penkert, H., Bergner, C. G., Su, M., Weil, M. T., Surma, M. A., Mann, M., Klose, C., & Simons, M. (2020). Cell-Type- and Brain-Region-Resolved Mouse Brain Lipidome. Cell Reports, 32(11), 108132. https://doi.org/10.1016/j.celrep.2020.108132

Freyre, C. A. C., Rauher, P. C., Ejsing, C. S., & Klemm, R. W. (2019). MIGA2 Links Mitochondria, the ER, and Lipid Droplets and Promotes De Novo Lipogenesis in Adipocytes. Molecular Cell, 76(5), 811–825. https://doi.org/10.1016/j.molcel.2019.09.011

Gao, Y., Vasilyev, D. V., Goncalves, M. B., Howell, F. V., Hobbs, C., Reisenberg, M., Shen, R., Zhang, M. Y., Strassle, B. W., Lu, P., Mark, L., Piesla, M. J., Deng, K., Kouranova, E. V., Ring, R. H., Whiteside, G. T., Bates, B., Walsh, F. S., Williams, G., … Doherty, P. (2010). Loss of retrograde endocannabinoid signaling and reduced adult neurogenesis in diacylglycerol lipase knock-out mice. Journal of Neuroscience, 30(6), 2017–2024. https://doi.org/10.1523/JNEUROSCI.5693-09.2010

Gehre, M., Bunina, D., Sidoli, S., Lübke, M. J., Diaz, N., Trovato, M., Garcia, B. A., Zaugg, J. B., & Noh, K. M. (2020). Lysine 4 of histone H3.3 is required for embryonic stem cell differentiation, histone enrichment at regulatory regions and transcription accuracy. Nature Genetics, 52(3), 273–282. https://doi.org/10.1038/s41588-020-0586-5

Haendel, M. A., Bollinger, K. E., & Baas, P. W. (1996). Cytoskeletal changes during neurogenesis in cultures of avian neural crest cells. Journal of Neurocytology, 25(1), 289–301. https://doi.org/10.1007/bf02284803

Han, X., Holtzman, D. M., & McKeel, D. W. (2001). Plasmalogen deficiency in early Alzheimer’s disease subjects and in animal models: Molecular characterization using electrospray ionization mass spectrometry. Journal of Neurochemistry, 77(4), 1168–1180. https://doi.org/10.1046/j.1471-4159.2001.00332.x

Han, X., Holtzman, D. M., McKeel, D. W., Kelley, J., & Morris, J. C. (2002). Substantial sulfatide deficiency and ceramide elevation in very early Alzheimer’s disease: Potential role in disease pathogenesis. Journal of Neurochemistry, 82(4), 809–818. https://doi.org/10.1046/j.1471-4159.2002.00997.x

Harayama, T., Eto, M., Shindou, H., Kita, Y., Otsubo, E., Hishikawa, D., Ishii, S., Sakimura, K., Mishina, M., & Shimizu, T. (2014). Lysophospholipid acyltransferases mediate phosphatidylcholine diversification to achieve the physical properties required in vivo. Cell Metabolism, 20(2), 295–305. https://doi.org/10.1016/j.cmet.2014.05.019

Hicks, A. M., DeLong, C. J., Thomas, M. J., Samuel, M., & Cui, Z. (2006). Unique molecular signatures of glycerophospholipid species in different rat tissues analyzed by tandem mass spectrometry. Biochimica et Biophysica Acta (BBA) - Molecular and Cell Biology of Lipids, 1761(9), 1022–1029. https://doi.org/10.1016/J.BBALIP.2006.05.010

Imgrund, S., Hartmann, D., Farwanah, H., Eckhardt, M., Sandhoff, R., Degen, J., Gieselmann, V., Sandhoff, K., & Willecke, K. (2009). Adult ceramide synthase 2 (CERS2)-deficient mice exhibit myelin sheath defects, cerebellar degeneration, and hepatocarcinomas. Journal of Biological Chemistry, 284(48), 33549–33560. https://doi.org/10.1074/jbc.M109.031971

Ingólfsson, H. I., Carpenter, T. S., Bhatia, H., Bremer, P. T., Marrink, S. J., & Lightstone, F. C. (2017). Computational Lipidomics of the Neuronal Plasma Membrane. Biophysical Journal, 113(10), 2271–2280. https://doi.org/10.1016/j.bpj.2017.10.017

Innis, S. M. (2007). Dietary (n-3) fatty acids and brain development. Journal of Nutrition, 137(4), 855–859. https://doi.org/10.1093/jn/137.4.855

Izant, J. G., & McIntosh, J. R. (1980). Microtubule-associated proteins: a monoclonal antibody to MAP2 binds to differentiated neurons. Proceedings of the National Academy of Sciences of the United States of America, 77(8), 4741–4745. https://doi.org/10.1073/pnas.77.8.4741

Janssen, C. I. F., Zerbi, V., Mutsaers, M. P. C., de Jong, B. S. W., Wiesmann, M., Arnoldussen, I. A. C., Geenen, B., Heerschap, A., Muskiet, F. A. J., Jouni, Z. E., van Tol, E. A. F., Gross, G., Homberg, J. R., Berg, B. M., & Kiliaan, A. J. (2015). Impact of dietary n-3 polyunsaturated fatty acids on cognition, motor skills and hippocampal neurogenesis in developing C57BL/6J mice. Journal of Nutritional Biochemistry, 26(1), 24–35. https://doi.org/10.1016/j.jnutbio.2014.08.002

Katsetos, C. D., Legido, A., Perentes, E., & Mörk, S. J. (2003). Class III β-tubulin isotype: A key cytoskeletal protein at the crossroads of developmental neurobiology and tumor neuropathology. Journal of Child Neurology, 18(12), 851–866. https://doi.org/10.1177/088307380301801205

Kim, H. Y. (2007). Novel Metabolism of Docosahexaenoic Acid in Neural Cells. Journal of Biological Chemistry, 282(26), 18661–18665. https://doi.org/10.1074/JBC.R700015200

Lauwers, E., Goodchild, R., & Verstreken, P. (2016). Membrane Lipids in Presynaptic Function and Disease. Neuron, 90(1), 11–25. https://doi.org/10.1016/j.neuron.2016.02.033

Levental, K. R., Malmberg, E., Symons, J. L., Fan, Y. Y., Chapkin, R. S., Ernst, R., & Levental, I. (2020). Lipidomic and biophysical homeostasis of mammalian membranes counteracts dietary lipid perturbations to maintain cellular fitness. Nature Communications, 11(1), 1–13. https://doi.org/10.1038/s41467-020-15203-1

Levy, M., & Futerman, A. H. (2010). Mammalian ceramide synthases. IUBMB Life, 62(5), 347–356. https://doi.org/10.1002/iub.319

Lin, C. Y. (1978). Properties of the thioesterase component obtained by limited trypsinization of the fatty acid synthetase multienzyme complex. Journal of Biological Chemistry, 253(6), 1954–1962. https://doi.org/10.1016/s0021-9258(19)62341-0

Metsalu, T., & Vilo, J. (2015). ClustVis: a web tool for visualizing clustering ofmultivariate data using Principal Component Analysisand heatmap. Nucleic Acids Research, 43(W1), W566–W570. https://doi.org/10.1093/nar/gkv468

Missmer, S. A., Suarez, L., Felkner, M., Wang, E., Merrill, A. H., Rothman, K. J., & Hendricks, K. A. (2006). Exposure to fumonisins and the occurence of neutral tube defects along the Texas-Mexico border. Environmental Health Perspectives, 114(2), 237–241. https://doi.org/10.1289/ehp.8221

Moore, S. A., Yoder, E., Murphy, S., Dutton, G. R., & Spector, A. A. (1991). Astrocytes, Not Neurons, Produce Docosahexaenoic Acid (22:6ω-3) and Arachidonic Acid (20:4ω-6). Journal of Neurochemistry, 56(2), 518–524. https://doi.org/10.1111/j.1471-4159.1991.tb08180.x

O’Brien, J. S., Fillerup, D. L., & Mead, J. F. (1964). Quantification and fatty acid and fatty aldehyde composition of ethanolamine, choline, and serine glycerophosphatides in human cerebral grey and white matter. Journal of Lipid Research, 5(3), 329–338. https://doi.org/10.1016/S0022-2275(20)40201-9

O’Brien, J. S., & Sampson, E. L. (1965). Fatty acid and fatty aldehyde composition of the major brain lipids in normal human gray matter, white matter, and myelin. Journal of Lipid Research, 6(4), 545–551. https://doi.org/10.1016/s0022-2275(20)39620-6

Pauling, J. K., Hermansson, M., Hartler, J., Christiansen, K., Gallego, S. F., Peng, B., Ahrends, R., & Ejsing, C. S. (2017). Proposal for a common nomenclature for fragment ions in mass spectra of lipids. PLoS ONE, 12(11), 1–21. https://doi.org/10.1371/journal.pone.0188394

Pfrieger, F. W., & Ungerer, N. (2011). Cholesterol metabolism in neurons and astrocytes. Progress in Lipid Research, 50(4), 357–371. https://doi.org/10.1016/j.plipres.2011.06.002

Pinot, M., Vanni, S., Pagnotta, S., Lacas-Gervais, S., Payet, L. A., Ferreira, T., Gautier, R., Goud, B., Antonny, B., & Barelli, H. (2014). Polyunsaturated phospholipids facilitate membrane deformation and fission by endocytic proteins. Science, 345(6197), 693–697. https://doi.org/10.1126/science.1255288

Puchkov, D., & Haucke, V. (2013). Greasing the synaptic vesicle cycle by membrane lipids. Trends in Cell Biology, 23(10), 493–503. https://doi.org/10.1016/j.tcb.2013.05.002

Salem, N., Litman, B., Kim, H. Y., & Gawrisch, K. (2001). Mechanisms of action of docosahexaenoic acid in the nervous system. Lipids, 36(9), 945–959. https://doi.org/10.1007/s11745-001-0805-6

Sampaio, J. L., Gerl, M. J., Klose, C., Ejsing, C. S., Beug, H., Simons, K., & Shevchenko, A. (2011). Membrane lipidome of an epithelial cell line. Proceedings of the National Academy of Sciences of the United States of America, 108(5), 1903–1907. https://doi.org/10.1073/pnas.1019267108

Sinensky, M. (1974). Homeoviscous adaptation: a homeostatic process that regulates the viscosity of membrane lipids in Escherichia coli. Proceedings of the National Academy of Sciences of the United States of America, 71(2), 522–525. https://doi.org/10.1073/pnas.71.2.522

Sladitschek, H. L., & Neveu, P. A. (2019). A gene regulatory network controls the balance between mesendoderm and ectoderm at pluripotency exit. Molecular Systems Biology, 15(12), 1–13. https://doi.org/10.15252/msb.20199043

Sprenger, R. R., Hermansson, M., Neess, D., Becciolini, L. S., Sørensen, S. B., Fagerberg, R., Ecker, J., Liebisch, G., Jensen, O. N., Vance, D. E., Færgeman, N. J., Klemm, R. W., & Ejsing, C. S. (2021). Lipid molecular timeline profiling reveals diurnal crosstalk between the liver and circulation. Cell Reports, 34(5), 108710. https://doi.org/10.1016/j.celrep.2021.108710

Stevens, V. L., & Tang, J. (1997). Fumonisin B1-induced sphingolipid depletion inhibits vitamin uptake via the glycosylphosphatidylinositol-anchored folate receptor. Journal of Biological Chemistry, 272(29), 18020–18025. https://doi.org/10.1074/jbc.272.29.18020

Swistowski, A., Peng, J., Han, Y., Swistowska, A. M., Rao, M. S., & Zeng, X. (2009). Xeno-free defined conditions for culture of human embryonic stem cells, neural stem cells and dopaminergic neurons derived from them. PLoS ONE, 4(7). https://doi.org/10.1371/journal.pone.0006233

Tu, J., Yin, Y., Xu, M., Wang, R., & Zhu, Z. J. (2018). Absolute quantitative lipidomics reveals lipidome-wide alterations in aging brain. Metabolomics, 14(1), 1–11. https://doi.org/10.1007/s11306-017-1304-x

Tulodziecka, K., Diaz-Rohrer, B. B., Farley, M. M., Chan, R. B., Paolo, G. Di, Levental, K. R., Neal Waxham, M., & Levental, I. (2016). Remodeling of the postsynaptic plasma membrane during neural development. Molecular Biology of the Cell, 27(22), 3480–3489. https://doi.org/10.1091/mbc.E16-06-0420

Valenza, M., Marullo, M., Di Paolo, E., Cesana, E., Zuccato, C., Biella, G., & Cattaneo, E. (2015). Disruption of astrocyte-neuron cholesterol cross talk affects neuronal function in Huntington’s disease. Cell Death and Differentiation, 22(4), 690–702. https://doi.org/10.1038/cdd.2014.162

Venkataraman, K., Riebeling, C., Bodennec, J., Riezman, H., Allegood, J. C., Cameron Sullards, M., Merrill, A. H., & Futerman, A. H. (2002). Upstream of growth and differentiation factor 1 (uog1), a mammalian homolog of the yeast longevity assurance gene 1 (LAG1), regulates N-stearoyl-sphinganine (C18-(dihydro)ceramide) synthesis in a fumonisin B1-independent manner in mammalian cells. Journal of Biological Chemistry, 277(38), 35642–35649. https://doi.org/10.1074/jbc.M205211200

Verheijen, M. H. G., Camargo, N., Verdier, V., Nadra, K., De Preux Charles, A. S., Médard, J. J., Luoma, A., Crowther, M., Inouye, H., Shimano, H., Chen, S., Brouwers, J. F., Helms, J. B., Feltri, M. L., Wrabetz, L., Kirschner, D., Chrast, R., & Smit, A. B. (2009). SCAP is required for timely and proper myelin membrane synthesis. Proceedings of the National Academy of Sciences of the United States of America, 106(50), 21383–21388. https://doi.org/10.1073/pnas.0905633106

Yamashita, A., Hayashi, Y., Nemoto-Sasaki, Y., Ito, M., Oka, S., Tanikawa, T., Waku, K., & Sugiura, T. (2014). Acyltransferases and transacylases that determine the fatty acid composition of glycerolipids and the metabolism of bioactive lipid mediators in mammalian cells and model organisms. Progress in Lipid Research, 53(1), 18–81. https://doi.org/10.1016/J.PLIPRES.2013.10.001

Ying, Q. L., Stavridis, M., Griffiths, D., Li, M., & Smith, A. (2003). Conversion of embryonic stem cells into neuroectodermal precursors in adherent monoculture. Nature Biotechnology, 21(2), 183–186. https://doi.org/10.1038/nbt780

Yousefi, S., Deng, R., Lanko, K., Salsench, E. M., Nikoncuk, A., van der Linde, H. C., Perenthaler, E., van Ham, T. J., Mulugeta, E., & Barakat, T. S. (2021). Comprehensive multi-omics integration identifies differentially active enhancers during human brain development with clinical relevance. Genome Medicine, 13(1), 1–27. https://doi.org/10.1186/s13073-021-00980-1

Zech, T., Ejsing, C. S., Gaus, K., De Wet, B., Shevchenko, A., Simons, K., & Harder, T. (2009). Accumulation of raft lipids in T-cell plasma membrane domains engaged in TCR signalling. EMBO Journal, 28(5), 466–476. https://doi.org/10.1038/emboj.2009.6

